## Supplementary File for "Target-agnostic identification of human antibodies to *Plasmodium falciparum* sexual forms reveals cross stage recognition of glutamate-rich repeats"

**This PDF file includes:**

Figs S1-9
Tables S1-4

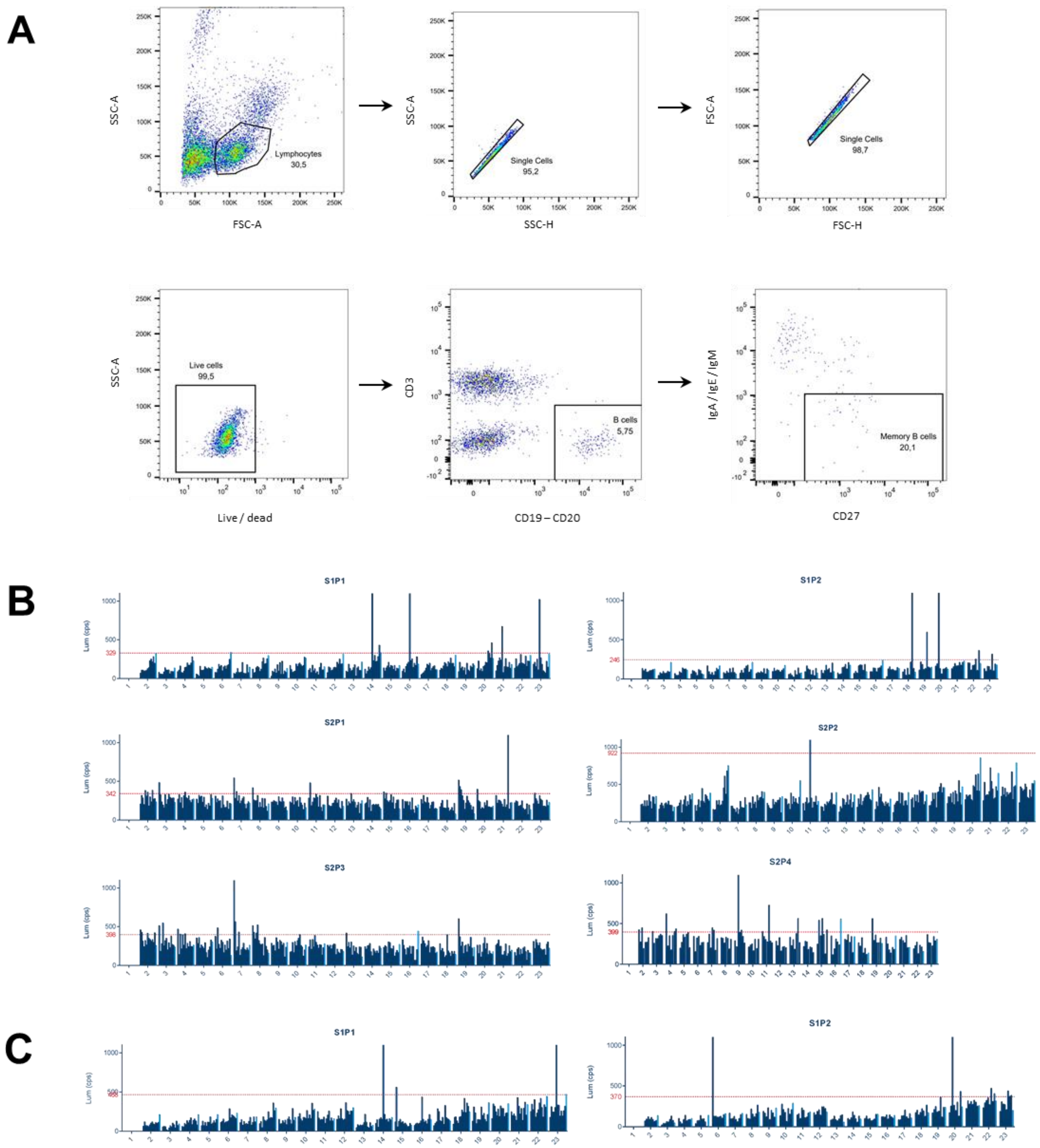


**Fig. S1. Memory B cell sorting and cell culture supernatant screening.** Gating strategy for agnostic MBCs sorting (A). Gamete extract (B) or gametocyte extract (C) ELISA for cell culture supernatant screening. Wells with signal above or close to positivity threshold (indicated in red) were selected for immunoglobulin variable genes amplification.

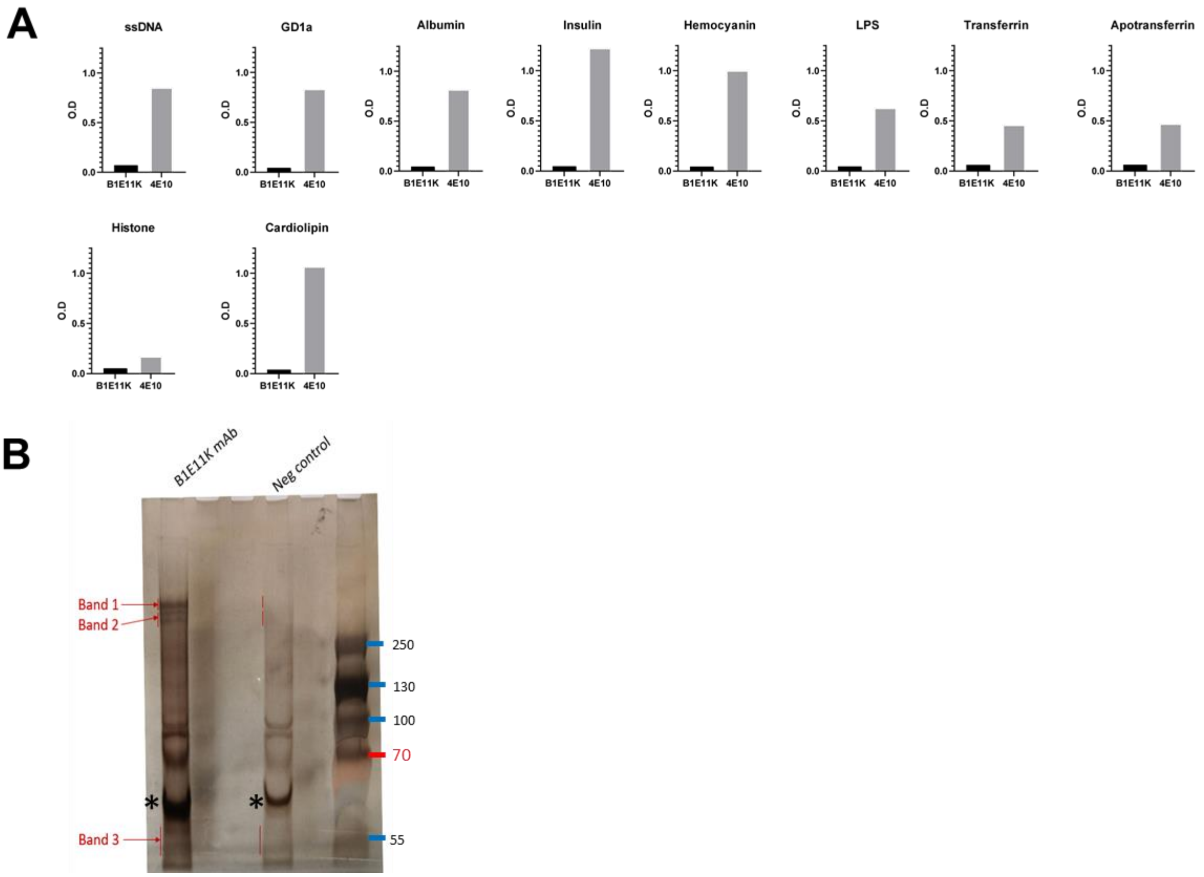


**Fig. S2. Further characterisation of B1E11K. (A)** B1E11K binding to a panel of human self-proteins, single-stranded DNA (ssDNA) and lipopolysaccharide (LPS) in ELISA. 4E10 is a polyreactive anti-HIV mAb (positive control). (B) Immunoprecipitation of B1E11K mAb against gametocyte extract. A Native Page 3-12% Bis Tris gel was used for protein separation followed by silver staining. VRC01, an anti-HIV mAb was used as a negative control. * : BSA.

977_PF3D7_0220000_e2s1_4 → Liver Stage Antigen 3

SDINKSWKKNTYVDKKLNKLFNRSLGESQVNGELASEEVKEKILDLLEEGNTLTESVDDNKNLEEAEDIKENILLSNIEEPKENIIDNLLNNIGQNSEKQESVSENVQVSDELFNELLNSVDVNGEVKENILEESQVNDDIF
NSLVKSVQQEQQHNVEEKVEESVEENDEESVEENVEENVEENDDESVASSVEESIASSVDESIDSSIEENVAPTVEEIVAPTVEEIVAPSVVESVAPSVEESVEENVEESVAENVEESVAENVEESVAENVEESVAENVEES
VAENVEESVAENVEEIVAPTVEESVAPTVEEIVAPSVEESVAPSVEEIVVPTVEESVAENVEEIVAPSVEEIVAPSVEEIVAPTVEESVAPTVEEIVAPSVEESVAPSVEEIVVPTVEESVAENVEESVAENVEEIVAPSVE
EIVAPSVEEIVAPSVEEIVAPSVEEIVAPSVEEIVAPSVEEIVAPSVEEIVAPSVEEIVAPTVEEIVAPTVEEIVAPSVEEIVAPTVEESVAENVATNLSDNLLSNLLGGIETEEIKDSILNEIEEVKENVVTTILENVEET
TAESVTTFSNILEEIQENTITNDTIEEKLEELHENVLSAALENTQSEEEKKEVIDVIEEVKEEVATTLIETVEQAEEESASTITEIFENLEENAVESNENVAENLEKLNETVFNTVLDKVEETVEISGESLENNEMDKAFFS
EIFDNVKGIQENLLTGMFRSIETSIVIQSEEKVDLNENVVSSILDN

1069_PF3D7_0309100_C_4 → OMD protein

MLEMNMIMKIYYIFLLFVFILIERCVDNDVINKTQENYLDHRDFEKDMDELDLSGLEYDIENVEEIAQKDSYNYVTQIYVYIRIDEEYKKYINHLELMRLGQRFMTILQNAMINVQLKKTGTGIFTCFYENKSITDDLATYF
LIQKEVDFIEVGFDKRYPEGRHAPITDSDMRVFIKNEELYPYDVP

113_PF3D7_1149200_e2s1_1 → ring infected erythrocyte surface antigen 3

GNFGYNGSSNSGAQFTDRCSRNLYCETLPINPYADSENPIVVSQVFGLPSEKPTFTLEGTPDIDHTNILGFNEKLMTDVNRYRYSNNYEAIPHTREFNPLIVDKVLFDYNEKVDNLGRSGGDIIKKMQTLWDEIMDINKRKY
DFLKTKLQKTYSQYKVQYDMPKEVYESKWGQCLKLINQGGDNLEERLNTQFKNWYRQKYLNLEEYRRLTVLNQIAWKALSNQIQYTCRKIMNSNISSFKHISELKSLEQRAAKDAQEEMRKRAEKQKKKKSKRRGWLCCGGG
DNETVEPQQEEPVQDVGEHQINEYGDILPSLKVSINNSAINYYDAVKDGKYLDDDSSDALYTDEDLLFDLEKQKYMDMLDGSEDESVEDNEEEHSGEANEEELSVDENVEEQNVDESGEQQSDDESGEHQSVNEIVEEQSVN
EIVEEQTVDEIVEQETVDENVEEQAVDENEEQQTVDENVEQQTIDESQVQEEISTIQENIEEVVSEVQQDSEVDRTLHVPDTRFYDILGVGVNADMK

637_PF3D7_0102200_e2s2_3 → ring infected erythrocyte surface antigen

NEAYQVLGDIDKKRWYNKYGYDGIKQVNFMNPSIFYLLSSLEKFKDFTGTPQIVTLLRFFFEKRLSMNDLENKSEHLLKFMEQYQKEREAHVSEYLLNILQPCIAGDSKWNVPIITKLEGLKGSRFDIPILESLRWIFKHVA
KTHLKKSSKSAKKLQQRTQANKQELANINNNLMSTLKEYVGSSEQMNSITYNFENINSNVDNGNQSKNISDLSYTDQKEILEKIVSYIVDISLYDIENTALNAAEQLLSDNSVDEKTLKKRAQSLKKLSSIMERYAGGKRND
KKAKKYDTQDVVGYIMHGISTINKEMKNQNENVPEHVQHNAEANVEHDAEENVEHDAEENVEENVEENVEENVEENVEENVEENVEENVEENVEENVEENVEENVEENVEENVEEYDEENVEEVEENVEEYDEENVEE
VEENVEENVEENVEENVEEYDEENVEEVEENVEENVEENVEENVEENVEEVEENVEENVEENVEENVEENVEENVEEYDEENVEEHNEEYDE

224_PF3D7_1127500_C_1 → Protein disulfide isomerase

MLEMIIKFLIFNLLLYLFRCNTKVNFTYPQNDFLRIDSENIEEAFENNDYWLINFYAPQCVYCKFIWDKLRYIQRSIQNENRTKIYFGEVNCDNQKGYTVCEIYGAMRVPQLKLFKGSELISTYSNDITNYNNIKSWIYYIT
TPLFVGVQDENELKSYETENNIFLTCSENLPADLFDVAKLYKEECYFINITNPDMCTRMNIKTNELHVMSAYNHSTYDLDTLDRELLKLFVNKNRFPLVMKIDHRNFFNIRSSGNNLMLLLLDMKKNPNAYILEYKQYAEKY
KNHNDILFGYIDGKYYEENLELYGTDSDYYPQIIVFSKNPREYYFEGYFNIDNVDNIIYDVKNKRIYHKFEEFTKFRIILIKFKKHIKYILKKAFKTDFLSFIGFICSVIMMLLTAILVIHTIYKYYMSYTHNTYIEKEEKS
EYPYDVP

236_PF3D7_1036300-S2_1 → Duffy binding like merozoite surface protein 2

INYTDIKTLSPSEYLIEKCPECKCTKKNLQDVFELTFDGKALLEKLKKEESPVSNSVNALPEPGQITLPDPSLKQTTQQENQPVVETPVTTAVINEHQGQTEPNKGDNNNERENHESNVGSIQEVNQGSVSEESHSKTIDPS
KIDDRLELSSGSSSLEQHSKEDVKKGCALELVPLSLSDIEQIANESEDVLEEIEEEINTDGEIEYITEEEIKEDIEEETEEDIEEETEEETEEETEEEADEETVKEIEDKPEQEIKNKSLEEKQIDKNTDTSEKKGFNNSEK
DEKARNLISKNYKNYNELDKNVHTLVNSIISLLEEGNGSDSTLNSLSKDITNLFKN

120_PF3D7_0411700_1 → conserved plasmodium protein

MKGLKVFMRSFVYFVLVCNLINNVIICHKINYENIEENPQINDQNIMEILNSSVLNNNMKKEKEKNILYDISYYIEYLKIINSLDNKYNDKFHKAILNIDTTKKKKEKKNIDLIYYSGKKYNNNIKDIKKQLHKSFYV

826_PF3D7_1038400_e2s1_3 → Pf 11.1

VIPEEVVEEIITEEIAEEIIPKELVEEVIPEEVVVEIIPEKLVEEFVSEELVEELVPEEIVEKVVPEELVEEIVPEELVEEVVPEELVEEVIPEELVEEFIPEEVVEEVLPEQLVEEVISEVVEEVVPEELVEEVIPEEVVE
ELVAEEVIEEVIPKELIEEAITEEVEEVIPEEIVEEVIPEEVVKEVIPEVEIEEILPEELIEELVAKEVIEEVITEEVAEEVIPEEIVQEVIPEEVVEEIITEVEIEEVLPEELIDEVVPEEVTEEVITEEVMPEEVVKEVI
PEEIAEEVIPKELVEEVIPEEVVEEIIPEVEIEEVLPEEVIEEVVPEVEIEELIPDEVVEEVLPEELVEEVIPEEVAVEVVPEEVIEEAVPEEIVEEFIPEEQIEEVIQEEIIEQVVPEELIEEVVPEEIIEEVIPEEIVEE
VIYEEVIPEELVEEVIAEKLVKEIVPEQVREEVTLEEIVEEMIPEEFVEEVAPEVEIEEIIPEELIEEVIPEVLVEEAVPEELIEKVIPEVLVEEIIYKEGIPEVLIEEVIPEELVQEVIPEEIVEEVVPEEFIEEMIPQEV
VKEVITEKFIEEMIPEEVVEEVIPEEFIEEMIPEVVVEEVVPEELVEEMLPEELIEVVVPEELIEEVIPEELVEEVVAEELVEEVIPEEL

402_PF3D7_1149200_e2s2_2 → ring infected erythrocyte surface antigen 3

PDTRFYDILGVGVNADMKEISESYFKLAKQYYPPKYSVNEGMLKFKQISEAYQILGDIDKRKMYNKFGYDGIKGVNFIHPTIYYMLASLEKFAFYTGSPQIVTLMKFLFEKKLTVNDLDTKSEHLSKIMGVYQKERETYISE
NLISRLQPYIDSIRNWDVQIKDQIYELMGSPFDIAIIDSIGWTLQYVSMSHMKNPKKAIKKLETRSKKNKETVAYENNKLMNILREYFGNNEQINSITYNMEYNTLNENNENGYRKILNLNHKKQKKLFEEIISYIVNISLS
DIENTVKNSAESILTVEGLDEKKLSKRIESLRMLANAIRKYILRGKKGKKYKNKDAKSLSGNIANEINLINKELQNLKEHTQANIPEHIEENVQENMEENVEENVEENVEENVEENVEENVEENVEENVEENVEENVEENIE
ENVEENVEENIEENIEENAEENVEENIEENIEENIEENIEENIEENVEENVEENIEENVEENVEENAEENAEENAEENAEENDETPQEHNEEYDE

696_PF3D7_0804500_e3_3 → uncharacterized protein

LKIYEILPGLQVVFIKEIEIKHDVSPKKCICEEKIHELKERRKKEEHENEGISIFNLSYFLSSNYHKKQSNTRSDEKMNSSYNYSYNRMPSDNNKSNNLLNGSAQNVNTNDSKKTVQLYSFQKNDKKKEKNTNSSFNSISHL
KSPSENSEEMDIHSDKEGYQSVIIGGINFGYLGFNFSKFMNNKKKITIIDAYLLSTTHLVTISNNGTIALIKI

651_PF3D7_1038400_e7s1_3 → Pf 11.1

KVVPEELVEEVIPEEIVEEVIPEELIEEFVPEELVEEEVPEELVEEVIPEVLVEEVIPEEFVEEVIPEELVEEVIPEELFEEVVPEELVEEVIPEEIVEEVIPEELIEEVVPEVLVEEVIPEELVEEVIPEELFEEVVPEEL
VAEVIPEEIVEEVIPEELIEEFVPEELVEEEVPEELVEEVIPEVLVEEEVPEELVEEVIPEELVEEMLPEELIEVVVPEELVEEVVPEIVEEVVPEELVEEVIPEEVVEEVLPEKLVEEVVPEIVEEVAPEVVEEVIPEELV
EEIIPEILVEEVVPIEVVEEAVPEVVVEEVVPEKIVEEVVPEKVLEEVIPELLVEEIVPIEVVGEIVPEEVVEKIVPMEVVEELLPEVVEEVPHEIVEEEVVKEVIPEDEKKVEIVEIIIEEKVEKIPKKKVIEKKKEKEKI
RKPKKKVEKEIKPKEVKGVKGKLKKDVKAKHLKGSKVKITEDIQKKMEDDIKEKIMEKELKEEIKDKEYLRKVIPKEEKKEPIPKKGKSVEPKKEKQAGVKKKEKVPETKKEKAPAPKKEKVQAPKKEKVPAPKKEKPAVPKK
EKVPEPKKEKVPEPKKEKAPAPKKEKVQAPKKEKAPAPKKEKAPAPKKEKAPAPKKEKAPAPKKEKPAVPKKEKPAVPKKEKVPETKKEKVPETKKEKVPETKKEKVPVPKKEKVPAPKKEKVPAPKKEKVPAPKKEKPAVP
KKEKVPETKKEKAPEPKKEKAPEPKKEKAPEPKKEKAPEPKKEKAPAPKKEKVPVPKKEKVPAPKKEKVPAPKKEKPAVPKKEKVPETKKEKEKDSQPKKEKAKETSRPPKKEKEKGK

643_PF3D7_1038400_e5s3_3 → Pf 11.1

EELIEVVVPEELVEEVVPEELVEEVVPEEVVEEVIPEELIEELVPEVEIEEILPEELIEELVPEEIIEEVIPKELIEEVIPEVVVEEVVPEVLVEEVVPEELVEEVIPEVLVEEEVPEELVEDVIPEELVEEVVPEELVEEV
IPEEVVEEVLPEELVEEVVPEEIVEEVIPEELVEEEVPEELVEEVIPEEVVEEVIPEEVPEKVIPEKLVEELIPEEVVEELIPEEVVEEMVPIEVVEEVVPTEVLEEVIPEIVVEEVVPEELVEEVIPEELVEEVIPEIVVE
EVVPEELVEEVIPEELVEEVVPEELVEEVVPEEIVEELLPKEIVEEVIYEEQVEEVIHEELVAEVIPEEVVEEVVPEVVEEVVPEIVAEVIPEEVVEEVVPEVVEEVVPEVVEEVIPEEVVEEVIPEEFIEEMIPEEIVEEV
VPEVVEEVIPEELVAEVIPEELVEEVIPEVLFEEEIPEVVVEEVVPEVVEEEIPEVVVEEVVPEVVEEVIPEEVVEELIPEEVVEEMVPIEIVEEVVPTEVLEEVIPEIVVEEVVPEELVEEVIPEELVEEVVPEELVEEVV
PEVVEEVVPEVVEEVVPEELVEEVIPEVVVEEVVPEELVEEVIPEELVEEIIPEEIVEEVIPEELIEEVVPEVLVEEVIPEELVEEVIPEELFEEVVPEELVAEVIPEEIVEEVIPEELIEEFVPEELVEEEVPEELVEEVI
PEVLVEEVIPEEFVEEVIPEEIVEEVIPEEIVEEVIPEEVVEEVVPEIVAEVIPEEVVEEVVPEIVAE

1070_PF3D7_0909000-S2_C_4 → uncharacterized protein

MLEYNDICVCINNLNNMKVKKAFFMVQDLFIKKYNCNKKKRKLFLRNIFISEILYRYKIIKKTEIIFNMNNEPPCDNYMKYSIISKRLYRNYTFDYIIDLIDVNGKVCFMKYISCLNKINHYFKNIKLKSIEIINSSSLENN
YSSFETLNSISSEEFSDLEKKNKNIYSKNNIKGRSKNQKEKKKKKKKSESMCDKCISNVYIDFSHFNLAYEDEDNEVFYYVTKKEYALLRRSSSFSYYIEKEGKKEENKTIKKKTVERVEDINMTHLDVEHDHIKEDDRKNS
EPEQYKKINKNKKNHLYINKKSHSIEQNSQNISLEKCDEKSDFNTVIYFYVNLLIPSNFIYIIASLIEATLFKSWIPFYTFPFKFGITDCKLLKERGLIDKMVYTKIAMPWIIRDRYLLMDVWICEDFEYSRGIFLYATNFP
RNTEINNNVNFDTDNCIEIDMSIHAFICPKSFDKTNVKCYVEISPNMNFNEFFISFITKVFIKSCITHFIKVCKEFEINEDYNKELQKNIFFYDKLRKAAKEYKIYYPYDVP

86_PF3D7_1038400_e1s1_1 → Pf 11.1

MKTKYKTLMVNIDDNDLGKRGNKIKPKHKKVEINLNDNNLSNNESKRKSRRKKVKVNLKDNLAKREGKMKRRNKKEKRRKEKDNKKMNELKLKREYKESYLLDNKSDMDVIRNDKFSYFTLYKNNLFPNSNNYFLHMLKTYD
EENKGIDENTDINENVEENEYVDKNTEGDKDGDEDKGGGESEGGDGHGSAGDSAGGDRGGDENKDKDNEENKDKKGNKDDDDEEEEEEEEEDDEEKEEKDEKDEKDEKDEKDEKDEKDEKDEKDEKDEKDEKDEKDEKGEKDE
SKGEEKFLKEEKMPKKEENMQDKIERDKEFEGFKDEYFTQMEGGAYGITVEHEGQKLRETLTEGVEGYEKVIEEMEQTKLDEFLLPTNEGKSDDFVSEMEEERLFNFVSYMEEEVKDKMFIEREQGTEEEVIHEDLSDEKIG
IEKKELETFVQDVSDEVLQDEQLIEEEKLTEEEKLTEEEKLTEEEKLTEDEQLIEEEKLTEEEKLTEEEKLTEEEKLTEEEKLTEEEKLTEEEKLTEEEKLTEEEKLTEEEIFIEEDKISPKEVEEPLEDVLMEKMEKEYKE
ILHEGKSVVTQDMDEKLLSENLEDDLIPEEIIEELVDEELVEEVTAEEIVEEFVAEVEIEEKIPKELVEEVMPEEDVDEVVPEELVEEVVPEELIEGVIPEELVEDVIPEKVLEKVIPEEIIEELVLEEVVEEIKHEELVEE
EIPEEVTEEVIPDEVVEEVIPEELIEKVVKEELVEKVVAEEIVEEVVPEELVEKVVAEEIVEEVIPEELIEEVITEKPAEEVIPEELVEEVIPKEIVEEVLPEEVVKEVIPEEVVEEVIPEELVEEVIPEEIVEEVVPEELV
EEMKPEEVVEEVISEELVEEMKPEEVVEEMKPEEVVEEVTPEKVVEEVVPEELVEEMKPEEVVEEVISEE

101_PF3D7_0522400_e3s1_1 → uncharacterized protein

YEMFLTAIEKLFSESTNSLKEPLKIHLKNSQSKILHKTRNGKYCSFSDYEMNERIKVGSLRKKKRKKEIDIMKKKEKIKNIYKKGYSDGSSFKKEERYYFDIPKNNTGFRITPKEGIELHVIFKKHDIKALNELKEDIKYFK
NVEMCILRHNKEYSSEHHLNNTPRDEMYVKKSCNCYNISKQVSMDENNKYNINIEHNDNNIENFENNNSNSNSNSNNYYMNNNFPVYNDPKQHNVNNINNEIISIDYDLIIENGFKNDVPPTFNNISNYENLEINFTSECFN
AFFNFIKYIEFYYLYKGGCTKIFNIVPSIEECIKYSEYRKTFNQKNSKMNDEYAEEFMNKFEASCPIHLLTKLRLLSDERIYSKEYFNNIQKGEWKKLFYNFIKNKKGKNYMIDENLEKLDRWFTHDMLFNLVLPQINFTYI
PLNLNHYNNNKIYYNKCFLSYNISCSFLFKQEKLSSMKKRYAIYMHNFNIHSLFVLFKRNKKFLINKKFCPIYLIEKLGIKNHCVYNEISHKCMNPYDIHPHNFNEEYLLEQNFITQNIQNGHNYNKDINDKNYKDNFNQNY
DYEKSKKNYKGSILNSLGKNTNNIDIHHEVNNNLPSDIMRSNSTCSNYKIHYEHNYNNIEHYDNTVNYKENKKCSHIITTKEKEAYMRSAGQRGRNHLFYNEYYNNTTKGRYKNMYMNNLNYNKYNDNNCDNNCDNNCDHYC
DNNCDNNNYYYEKDYMCHIFFNNDNKVNDNMQYKYIDGNIDTNNVDKKKIPIENLQSFKNEYISQKHNMNKNKMIHENDINYNKNNVDSKTDDKKGFWKFILGKKYKGEDDSIKNNKSNYNTILGTNFDLLIYNKINKLPNC
TMIFINDYSNNIREMDIYICKYIININNLIPIMNFIPKNNICLEIFLKYDNIIDKDVIEALTLYLSMGSARKKIVQVCIEQL

**Fig. S3. Sequences of the recombinant protein domains used in the microarray.**

## A

>tr|Q8I0U6|Q8I0U6_PLAF7 Ring-infected erythrocyte surface antigen OS=Plasmodium falciparum (isolate 3D7) OX=36329 GN=PF3D7_0102200 PE=4 SV=1

MRPFHAYSWIFSQQYMDTKNVKEKNPTIYSFDDEEKRNENKSFLKVLCSKRGVLPIIGILYIILNGNLGYNGSSSSGVQFTDRCSRNLYGETLPVNPYADSENPIVVSQVFGLPFEK
PTFTLESPPDIDHTNILGFNEKFMTDVNRYRYSNNYEAIPHISEFNPLIVDKVLFDYNEKVDNLGRSGGDIIKKMQTLWDEIMDINKRKYDSLKEKLQKTYSQYKVQYDMPKEAYESK
WTQCIKLIDQGGENLEERLNSQFKNWYRQKYLNLEEYRRLTVLNQIAWKALSNQIQYSCRKIMNSDISSFKHINELKSLEHRAAKAAEAEMKKRAQKPKKKKSRRGWLCCGGGDI
ETVEPQQEEPVQTVQEQQVNEYGDILPSLRASITNSAINYYDTVKDGVYLDHETSDALYTDEDLLFDLEKQKYMDMLDTSEEESVKENEEEHTVDDEHVEEHTADDEHVEEPTVA
DDEHVEEPTVADEHVEEPTVAEEHVEEPTVAEEHVEEPASDVQQTSEAAPTIEIPDTLYYDILGVGVNADMNEITERYFKLAENYYPYQRSGSTVFHNFRKVNEAYQVLGDIDKKR
WYNKYGYDGIKQVNFMNPSIFYLLSSLEKFKDFTGTPQIVTLLRFFFEKRLSMNDLENKSEHLLKFMEQYQKEREAHVSEYLLNILQPCIAGDSKWNVPIITKLEGLKGSRFDIPILES
LRWIFKHVAKTHLKKSSKSAKKLQQRTQANKQELANINNNLMSTLKEYVGSSEQMNSITYNFENINSNVDNGNQSKNISDLSYTDQKEILEKIVSYIVDISLYDIENTALNAAEQLLSD
NSVDEKTLKKRAQSLKKLSSIMERYAGGKRNDKKAKKYDTQDVVGYIMHGISTINKEMKNQNENVPEHVQHNAEANVEHAEENVEHDAEENVEENVEENVEENVEENVEENVEE
NVEENVEENVEENVEENVEENVEENVEENVEENVEEYDEENVEEVEENVEEYDEENVEEVEENVEENVEENVEENVEEYDEENVEEVEENVEENVEENVEENVEENVEEVEEN
VEENVEENVEENVEENVEENVEEYDEENVEEHNEEYDE

## B

>tr|Q8IHN1|Q8IHN1_PLAF7 Ring-infected erythrocyte surface antigen OS=Plasmodium falciparum (isolate 3D7) OX=36329 GN=PF3D7_1149200 PE=4 SV=1

MKPYSSYSSAFSKQYMGTKSVKAKNPTIYSFEEEKQNENMSLLKSLCSKRLVLPILGILYIILNGNFGYNGSSNSGAQFTDRCSRNLYCETLPINPYADSENPIVVSQVFGLPSEKPTF
TLEGTPDIDHTNILGFNEKLMTDVNRYRYSNNYEAIPHTREFNPLIVDKVLFDYNEKVDNLGRSGGDIIKKMQTLWDEIMDINKRKYDFLKTKLQKTYSQYKVQYDMPKEVYESKWG
QCLKLINQGGDNLEERLNTQFKNWYRQKYLNLEEYRRLTVLNQIAWKALSNQIQYTCRKIMNSNISSFKHISELKSLEQRAAKDAQEEMRKRAEKQKKKKSKRRGWLCCGGGDNE
TVEPQQEEPVQDVGEHQINEYGDILPSLKVSINNSAINYYDAVKDGKYLDDDSSDALYTDEDLLFDLEKQKYMDMLDGSEDESVEDNEEEHSGEANEEELSVDENVEEQNVDESG
EQQSDDESGEHQSVNEIVEEQSVNEIVEEQTVDEIVEQETVDENVEEQAVDENEEQQTVDENVEQQTIDESQVQEEISTIQENIEEVVSEVQQDSEVDRTLHVPDTRFYDILGVGV
NADMKEISESYFKLAKQYYPPKYSVNEGMLKFKQISEAYQILGDIDKRKMYNKFGYDGIKGVNFIHPTIYYMLASLEKFAFYTGSPQIVTLMKFLFEKKLTVNDLDTKSEHLSKIMGVY
QKERETYISENLISRLQPYIDSIRNWDVQIKDQIYELMGSPFDIAIIDSIGWTLQYVSMSHMKNPKKAIKKLETRSKKNKETVAYENNKLMNILREYFGNNEQINSITYNMEYNTLNENN
ENGYRKILNLNHKKQKKLFEEIISYIVNISLSDIENTVKNSAESILTVEGLDEKKLSKRIESLRMLANAIRKYILRGKKGKKYKNKDAKSLSGNIANEINLINKELQNLKEHTQANIPEHIEEN
VQENMEENVEENVEENVEENVEENVEENVEENVEENVEENVEENVEENIEENVEENVEENIEENIEENAEENVEENIEENIEENIEENIEENIEENVEENVEENIEENVEENVEENAEE
NAEENAEENAEENDETPQEHNEEYDE

## C

>tr|O96275|O96275_PLAF7 Liver stage antigen 3 OS=Plasmodium falciparum (isolate 3D7) OX=36329 GN=PF3D7_0220000 PE=4 SV=1

MTNSNYKSNNKTYNENNNEQITTIFNRTNMNPIKKCHMREKINKYFFLIKILTCTILIWAVQYANNSDINKSWKKNTYVDKKLNKLFNRSLGESQVNGELASEEVKEKILDLLEEGNTLT
ESVDDNKNLEEAEDIKENILLSNIEEPKENIIDNLLNNIGQNSEKQESVSENVQVSDELFNELLNSVDVNGEVKENILEESQVNDDIFNSLVKSVQQEQQHNVEEKVEESVEENDEESV
EENVEENVEENDDESVASSVEESIASSVDESIDSSIEENVAPTVEEIVAPTVEEIVAPSVVESVAPSVEESVEENVEESVAENVEESVAENVEESVAENVEESVAENVEESVAENVEE
SVAENVEEIVAPTVEESVAPTVEEIVAPSVEESVAPSVEEIVVPTVEESVAENVEEIVAPSVEEIVAPSVEEIVAPTVEESVAPTVEEIVAPSVEESVAPSVEEIVVPTVEESVAENVEES
VAENVEEIVAPSVEEIVAPSVEEIVAPSVEEIVAPSVEEIVAPSVEEIVAPSVEEIVAPSVEEIVAPSVEEIVAPTVEEIVAPTVEEIVAPSVEEIVAPTVEESVAENVATNLSDNLLSNLLG
GIETEEIKDSILNEIEEVKENVVTTILENVEETTAESVTTFSNILEEIQENTITNDTIEEKLEELHENVLSAALENTQSEEEKKEVIDVIEEVKEEVATTLIETVEQAEEESASTITEIFENLEE
NAVESNENVAENLEKLNETVFNTVLDKVEETVEISGESLENNEMDKAFFSEIFDNVKGIQENLLTGMFRSIETSIVIQSEEKVDLNENVVSSILDNIENMKEGLLNKLENISSTEGVQET
VTEHVEQNVYVDVDVPAMKDQFLGILNEAGGLKEMFFNLEDVFKSESDVITVEEIKDEPVQKEVEKETVSIIEEMEENIVDVLEEEKEDLTDKMIDAVEESIEISSDSKEETESIKDKEK
DVSLVVEEVQDNDMDESVEKVLELKNMEEELMKDAVEINDITSKLIEETQELNEVEADLIKDMEKLKELEKALSEDSKEIIDAKDDTLEKVIEEEHDITTTLDEVVELKDVEEDKIEKVSD
LKDLEEDILKEVKEIKELESEILEDYKELKTIETDILEEKKEIEKDHFEKFEEEAEEIKDLEADILKEVSSLEVEEEKKLEEVHELKEEVEHIISGDAHIKGLEEDDLEEVDDLKGSILDMLKG
DMELGDMDKESLEDVTAKLGERVESLKDVLSSALGMDEEQMKTRKKAQRPKLEEVLLKEEVKEEPKKKITKKKVRFDIKDKEPKDEIVEVEMKDEDIDEDIEEDVEEDIEEDKVEDID
EDIDEDIDEDIGEDKDEVIDLIVQKEKRIEKVKEKKKKLEKKVEEGVSGLKKHVDEVMKYVQKIDKEVDKEVSKALESKNDVTNVLKQNQDFFSKVKNFVKKYKVFAAPFISAVAAFAS
YVVGFFTFSLFSSCVTIASSTYLLSKVDKTINKNKERPFYSFVFDIFKNLKHYLQQMKEKFSKEKNNNVIEVTNKAEKKGNVQVTNKTEKTTKVDKNNKVPKKSRTQKSK

## D

>sp|P68875|P230_PLAFO Gametocyte surface protein P230 OS=Plasmodium falciparum (isolate NF54) OX=5843 GN=PF230 PE=1 SV=1

MKKIITLKNLFLIILVYIFSEKKDLRCNVIKGNNIKDDEDKRFHLFYYSHNLFKTPETKEKKNKKECFYKNGGIYNLSKEIRMRKDTSVKIKQRTCPFHKEGSSFEMGSKNITCFYPIVGK
KERKTLDTIIIKKNVTNDHVVSSDMHSNVQEKNMILIRNIDKENKNDIQNVEEKIQRDTYENKDYESDDTLIEWFDDNTNEENFLLTFLKRCLMKIFSSPKRKKTVVQKKHKSNFFINSS
LKYIYMYLTPSDSFNLVRRNRNLDEEDMSPRDNFVIDDEEEEEEEEEEEEEEEEEEEEEEEEEEYDDYVYEESGDETEEQLQEEHQEEVGAESSEESFNDEDEDSVEARDGDMIRV
DEYYEDQDGDTYDSTIKNEDVDEEVGEEVGEEVGEEVGEEVGEEVGEEVGEEVGEEVGEEEGEEVGEGVGEEVGEEEGEEVGEEEGEYVDEKERQGEIYPFGDEEEKDEGGE
SFTYEKSEVDKTDLFKFIEGGEGDDVYKVDGSKVLLDDDTISRVSKKHTARDGEYGEYGEAVEDGENVIKIIRSVLQSGALPSVGVDELDKIDLSYETTESGDTAVSEDSYDKYASNN
TNKEYVCDFTDQLKPTESGPKVKKCEVKVNEPLIKVKIICPLKGSVEKLYDNIEYVPKKSPYVVLTKEETKLKEKLLSKLIYGLLISPTVNEKENNFKEGVIEFTLPPVVHKATVFYFICDN
SKTEDDNKKGNRGIVEVYVEPYGNKINGCAFLDEDEEEEKYGNQIEEDEHNEKIKMKTFFTQNIYKKNNIYPCYMKLYSGDIGGILFPKNIKSTTCFEEMIPYNKEIKWNKENKSLGNL
VNNSVVYNKEMNAKYFNVQYVHIPTSYKDTLNLFCSIILKEEESNLISTSYLVYVSINEELNFSLFDFYESFVPIKKTIQVAQKNVNNKEHDYTCDFTDKLDKTVPSTANGKKLFICRKHL
KEFDTFTLKCNVNKTQYPNIEIFPKTLKDKKEVLKLDLDIQYQMFSKFFKFNTQNAKYLNLYPYYLIFPFNHIGKKELKNNPTYKNHKDVKYFEQSSVLSPLSSADSLGKLLNFLDTQET
VCLTEKIRYLNLSINELGSDNNTFSVTFQVPPYIDIKEPFYFMFGCNNNKGEGNIGIVELLISKQEEKIKGCNFHESKLDYFNENISSDTHECTLHAYENDIIGFNCLETTHPNEVEVEVE
DAEIYLQPENCFNNVYKGLNSVDITTILKNAQTYNINNKKTPTFLKIPPYNLLEDVEISCQCTIKQVVKKIKVIITKNDTVLLKREVQSESTLDDKIYKCEHENFINPRVNKTFDENVEYTC
NIKIENFFNYIQIFCPAKDLGIYKNIQMYYDIVKPTRVPQFKKFNNEELHKLIPNSEMLHKTKEMLILYNEEKVDLLHFYVFLPIYIKDIYEFNIVCDNSKTMWKNQLGGKVIYHITVSKRE
QKVKGCSFDNEHAHMFSYNKTNVKNCIIDAKPKDLIGFVCPSGTLKLTNCFKDAIVHTNLTNINGILYLKNNLANFTYKHQFNYMEIPALMDNDISFKCICVDLKKKKYNVKSPLGPKVL
RALYKKLNIKFDNYVTGTDQNKYLMTYMDLHLSHKRNYLKELFHDLGKKKPADTDANPESIIESLSINESNESGPFPTGDVDAEHLILEGYDTWESLYDEQLEEVIYNDIESLELKDIE
QYVLQVNLKAPKLMMSAQIHNNRHVCDFSKNNLIVPESLKKKEELGGNPVNIHCYALLKPLDTLYVKCPTSKDNYEAAKVNISENDNEYELQVISLIEKRFHNFETLESKKPGNGDVV
VHNGVVDTGPVLDNSTFEKYFKNIKIKPDKFFEKVINEYDDTEEEKDLESILPGAIVSPMKVLKKKDPFTSYAAFVVPPIVPKDLHFKVECNNTEYKDENQYISGYNGIIHIDISNSNRKI
NGCDFSTNNSSILTSSVKLVNGETKNCEININNNEVFGIICDNETNLDPEKCFHEIYSKDNKTVKKFREVIPNIDIFSLHNSNKKKVAYAKVPLDYINKLLFSCSCKTSHTNTIGTMKVTL
NKDEKEEEDFKTAQGIKHNNVHLCNFFDNPELTFDNNKIVLCKIDAELFSEVIIQLPIFGTKNVEEGVQNEEYKKFSLKPSLVFDDNNNDIKVIGKEKNEVSISLALKGVYGNRIFTFDKN
GKKGEGISFFIPPIKQDTDLKFIINETIDNSNIKQRGLIYIFVRKNVSENSFKLCDFTTGSTSLMELNSQVKEKKCTVKIKKGDIFGLKCPKGFAIFPQACFSNVLLEYYKSDYEDSEHINY
YIHKDKKYNLKPKDVIELMDENFRELQNIQQYTGISNITDVLHFKNFNLGNLPLNFKNHYSTAYAKVPDTFNSIINFSCNCYNPEKHVYGTMQVESDNRNFDNIKKNENVIKNFLLPNIE
KYALLLDDEERQKKIKQQQEEEQQEQILKDQDDRLSRHDDYNKNHTYILYDSNEHICDYEKNESLISTLPNDTKKIQKSICKINAKALDVVTIKCPHTKNFTPKDYFPNSSLITNDKKIVI
TFDKKNFVTYIDPTKKTFSLKDIYIQSFYGVSLDHLNQIKKIHEEWDDVHLFYPPHNVLHNVVLNNHIVNLSSALEGVLFMKSKVTGDETATKKNTTLPTDGVSSILIPPYVKEDITFHLF
CGKSTTKKPNKKNTSLALIHIHISSNRNIIHGCDFLYLENQTNDAISNNNNNSYSIFTHNKNTENNLICDISLIPKTVIGIKCPNKKLNPQTCFDEVYYVKQEDVPSKTITADKYNTFSKDKI
GNILKNAISINNPDEKDNTYTYLILPEKFEEELIDTKKVLACTCDNKYIIHMKIEKSTMDKIKIDEKKTIGKDICKYDVTTKVATCEIIDTIDSSVLKEHHTVHYSITLSRWDKLIIKYPTNEKT
HFENFFVNPFNLKDKVLYNYNKPINIEHILPGAITTDIYDTRTKIKQYILRIPPYVHKDIHFSLEFNNSLSLTKQNQNIIYGNVAKIFIHINQGYKEIHGCDFTGKYSHLFTYSKKPLPNDDDI
CNVTIGNNTFSGFACLSHFELKPNNCFSSVYDYNEANKVKKLFDLSTKVELDHIKQNTSGYTLSYIIFNKESTKLKFSCTCSSNYSNYTIRITFDPNYIIPEPQSRAIIKYVDLQDKNFAK
YLRKL

# E

>tr|Q8I6U6|Q8I6U6_PLAF7 Gametocyte-specific protein OS=Plasmodium falciparum (isolate 3D7) OX=36329
GN=PF3D7_1038400 PE=4 SV=2

MKKESYRFLNISINNGSDKKKKSNFFVKIKNGTDLKDTKKDRIRFHFYFYFFIVKIILLSLLIWTVIYSNNCSNGKYKSTLYIVGNHLRFRGFRILAENEYEMKTKYKTLMVNIDDNDLGKRG
NKIKPKHKKVEINLNDNNLSNNESKRKSRRKKVKVNLKDNLAKREGKMKRRNKKEKRRKEKDNKKMNELKLKREYKESYLLDNKSDMDVIRNDKFSYFTLYKNNLFPNSNNYFLHMLK
TYDEENKGIDENTDINENVEENEYVDKNTEGDKDGDEDKGGGESEGGDGHGSAGDSAGGDRGGDENKDKDNEENKDKKGNKDDDDEEEEEEEEDDEEKEEKDEKDEKDEKDEK
DEKDEKDEKDEKDEKDEKDEKDEKDEKDEKGEKDESKGEEKFLKEEKMPKKEENMQDKIERDKEFEGFKDEYFTQMEGGAYGITVEHEGQKLRETLTEGVEGYEKVIEEMEQTKLDEFLL
PTNEGKSDDFVSEMEEERLFNFVSYMEEEVKDKMFIEREQGTEEEVIHEDLSDEKIGIEKKELETFVQDVSDEVLQDEQLIEEEKLTEEEKLTEEEKLTEEEKLTEDEQLIEEEKLTEEEK
LTEEEKLTEEEKLTEEEKLTEEEKLTEEEKLTEEEKLTEEEKLTEEEIFIEEDKISPKEVEEPLEDVLMEKMEKEYKEILHEGKSVVTQDMDEKLLSENLEDDLIPEEIIEELVDEELVEEVTA
EEIVEEFVAEVEIEEKIPKELVEEVMPEEDVDEVVPEELVEEVVPEELIEGVIPEELVEDVIPEKVLEKVIPEEIIEELVLEEVVEEIKHEELVEEEIPEEVTEEVIPDEVVEEVIPEELIEKVVKE
ELVEKVVAEEIVEEVVPEELVEKVVAEEIVEEVIPEELIEEVITEKPAEEVIPEELVEEVIPKEIVEEVLPEEVVKEVIPEEVVEEVIPEELVEEVIPEEIVEEVVPEELVEEMKPEEVVEEVISE
ELVEEMKPEEVVEEMKPEEVVEEVTPEKVVEEVVPEELVEEMKPEEVVEEVISEELVEEMKPEEVVEEVTPEKVVEEVVPEELVEEMKPEEVVEEVISEELVEEMKPEEVVEEVISEEL
VEEMKPEEVVEEVTPEKVVEEVVPEELVEEMKPEEVVEEVISEELVEEMKPEEVVEEVTPEKVVEEVVPEELVEEMKPEEVVEEVISEELVEEMKPEEVVEEVVPEELVEEVIPEEVVE
EIIPEEVVHEAKHEELFEKTVPEVEVQEIIPKELVEVIPEEVLPEELIEEVIPEEIIEEVIPEEVVEEVIPEEIVQEVIPEDLMEEVVPEEIIEEKILEEIVEEEIHEEIVEEIISEEVVEEVVPAEVVE
EMIPEKLVEEVIPEELVEEVISEELVEEVVPVEVLEEVVPVEVLEEVIPEEVVEEVIPEEFIEEMIPEEIVEEVVPEVVEEVIPEELVEEVVPVELLEEVIPEELLEEVIPEELVEEVVPEELVED
VIPEELIAEVVSEELVEEEIPEEVLEEVIPEELVEEVIPEELVEEVIPEVLVEEVIPEELVEEVIPEVLVEEEVPEELVEEVIPELLVAEVIPEELVAEVIPEEFVEEVIPEEVVEELVPEELVEEVI
PEEIVEEVVEEVVEEIIPEEVIEEVIPEEIVKEVVPEVLVEEVVPEEIVEEVVPEEIVEEVVPEELVEEVVPEELIEEVIPEELVEEVIPEVLVEEVIPEVLVEEMVLEELVEEVIPEVLVEEMVP
EEVVEKMVLEEVVEEVIPEEVVEEVIPEEDVEQVIPEELIEEVVPEEVIEEVITEEVVEEVLPEELVEKVVEEEILEEVIPEELVEEVVREELIEEVVTEELVEKVVEEEIVEEVVLAEVVEEVI
PQEIVEEVIPEEVVEEVIPKEIVEEVIPEELVEEVVPEELVAEVIPEELVAEVIPEELIEEVVTEELVEEVIPEEVVEEVIPEEVVEEVVPEELVEEVVPEELVEEVVPEEIVEELLPKEVVEEVI
YEEQVEEVIPEELVAEVIPEKLVEEVIPEELVEEVIPEELVEEVVPEVVEEVVPEELVEEVIPEELVEEVIPEVLVEEEVPEELVEEVIPEELVEEMLPEELIEVVVPEELVEEVVPEELVEEV
VPEEVVEEVIPEELIEELVPEVEIEEILPEELIEELVPEEIIEEVIPKELIEEVIPEVVVEEVVPEVVEEVVPEIVEEVVPEVVEEVVPEEIIEEKISEEVVEEVLPKEIVEEVIPEELVEEVVPEELV
AEVIPEELVEEVIPEVLFEEEIPEVVVEEVVPEVVEEEIPEVVVEEVVPEVVEEVIPEELVEEVIPEELIEEVIPEEVAEEVLPEELIEEVIPEELVEEVVPEELVAEVIPEEIVEEVVREELVEE
VVPEEVVEEVIPKEIVEEVVSEEVVEEVIPKEIVEEEVPEELVEVVPEELVEEVVPEELVEEVVPEELVEEVVPEELVEEVVPEELVEEVVPEVVEEVQEQLVEEFIPEEVVEELVPEELVE
EVIPEELVEEVIPEEIVEEVLPEEVVKEVIPEEVVEEVIPEELVEEVIPEEIVEEVLPEEVVKEVIPEEVVEEVIPEELVEEVIPEELVEEVIPEELVEEVVSEELVEEVIPEEIVEEVIPEELLEEV
VEEVVPEIVEEVVPEVVEEVITEEVIPEELIKEVIPEELVEEVIPEEIVEEVIPEEIVEEVISEEVVEEVVPEVVVEEVIPEEIVEEVLPKEIVEEVISEKVVEEMIPEELVEEVIPEEVVEEVIPEV
EIEEILPEELIEELVAEEVIEEVIPKELIEEVIPEKLIEDVVPEEIVEEVLPEEVVKEVIPEEVVEEVLPEELVEEVIPEVVEEVVPEELVEEVLPKEVVEEVIPEVEIEEILPEEIIEELVAKEVIEEV
IPKELIEEAITEEVVEEMIPEEIIEEVVPEEVIEEVITEEVVEEVVPEEVIEEVITEEVVEEAIPEEVVEEVIPEVEIEEVLPEELIEELVPEEVTEEVITEEVVEEAISEVVEEVIPEEIVKEVIPEEV
VEEIITEEIAEEIIPKELVEEVIPEEVVVEIIPEKLVEEFVSEELVEELVPEEIVEKVVPEELVEEIVPEELVEEVVPEELVEEVIPEELVEEFIPEEVVEEVLPEQLVEEVISEVVEEVVPEELVE
EVIPEEVVEELVAEEVIEEVIPKELIEEAITEEVEEVIPEEIVEEVIPEEVVKEVIPEVEIEEILPEELIEELVAKEVIEEVITEEVAEEVIPEEIVQEVIPEEVVEEIITEVEIEEVLPEELIDEVVPEE
VTEEVITEEVMPEEVVKEVIPEEIAEEVIPKELVEEVIPEEVVEEIIPEVEIEEVLPEEVIEEVVPEVEIEELIPDEVVEEVLPEELVEEVIPEEVAVEVVPEEVIEEAVPEEIVEEFIPEEQIEEVI
QEEIIEQVVPEELIEEVVPEEIIEEVIPEEIVEEVIYEEVIPEELVEEVIAEKLVKEIVPEQVREEVTLEEIVEEMIPEEFVEEVAPEVEIEEIIPEELIEEVIPEVLVEEAVPEELIEKVIPEVLVEEII
YKEGIPEVLIEEVIPEELVQEVIPEEIVEEVVPEEFIEEMIPQEVVKEVITEKFIEEMIPEEVVEEVIPEEFIEEMIPEVVVEEVVPEELVEEMLPEELIEVVVPEELIEEVIPEELVEEVVAEELV
EEVIPEELVEKVVPEVVEEFIHEVVEEVVPEVVEEVVPEVVEEVVPEVVEEVVPEVVEEVVPEVVEEVVPEVVEEVVPEVVEEVVPEVVEEVVPEVVEEVVPEVVEEVVPEELVEEVIPEVLFEEEIPE
ELVEVVPVELLEEAIPEELVEEVIPEELVEEVIPEELVEEVIPEELVEEVILEELVEEVIPEELVEEVIPEELVEEVIPEVLVEEEIPEKLVEEDIPEKLVEEVVPEELVEVVIPEELVEEVIPEELV
KEVIPEELVKEVVPEELVEEEIPKVVEEVLPEELVEEVMLEEVVEEVVPEIVEEVVPEELVEEVVPEVVEEVVPEEIVEEVIPEELVEEVVKEELVEEVVKEELVEEVVKEELVEEVIPEVLV
EEVVPEELLEEVIPKVLVEEVIPELLVEEVVPEVIEEVVPEVVEEVVPEELLEEVIPEVLVEEEIPEKLVEEVVPEELVEEVIPEVLVEEVVPEELVEEVISEELVEEVIPEELVEEVIPEEVVE
EVVPEVVEEVVPEVVEEVVPEVVEEVVPEVVEEVVPEVVEEVVPEVVEEVVPEVVEEVVPEVVEEVVPEVVEEVVPEELIEEVIPEEIVEEVIPEELVAEVIPEKLVEEVIPEELVEEVIPEE
LVEEVIPEELVEEVLPEEVVEEAVPEVVEEIVPEELIEEVIPEELIKEVVPEEVIEEVITEEVVEEVLPEEIVEEVLPEEVIEEVVPEEVLPEEVVEEVVPEELVEEVVPEELVEEVIPEELVEEV
LPEEVVEEVVPKELVEELIPEELVEEVIPEELVEEVVPEELVAEVIPEELVEEVIPEELVEEVIPEELVEEEIPEELVEEVILEELVEEVVPEELIEEVVPEELIEEVVPEELIEEVVEKVIPEEIIE
EVVPEEVIEEVITEEVVEEVVPEKVIEEVITEEVVEEVVPEEVIDEVITEEVVEEVVPEEVIEEVIIEEVVEEVVPEEVIEEVITEEVVEEAIPEELVEEVIPEVLVEEEIPEELVEEVILEELVEEV
VAEELIEEVVEEVIPEKLVEEVIPEEIVEEVIPEELVEVVVPEELVEEVIPEELVEEVIHEELVEEVIPEELVEEVIHEEIVEEVIPEEVVEEVIPEEIVEEVVPEELLEEVIPELLVEEVVPEVVEE
VVPVELLEEVVEEVITEEVIPEELIKEVISEELVEELIPEEVVQEVITEEVLEEVIPEEIVEELIPEKLIGEVVPEKVVEEVVLEEVVEEVIPEEIVEEVITEEVLAEEIVEEFVPEELVEEMLPEEV
VEERVREVEIEEMVPEVLVEEVLPEVVEEVITEEVVEEMVPEEVVEEMVPKEIVEEVLPEEVVEEVIPEEVVEEMVPEVLVEEVVPEEIVEKVVPEEVVEEVVAEEIIEEVITEEVVEEIVPG
EVVEEMVPEEIIEEVVPEEVIEEVITEEVVEEVIPEELVEEVIPEISVEVEIPEVLVEEVIPEKLVKVVPEELVEEVIPEELVEEVITEALVEEVVDVVVPEEIFEEVLPEEVVEEVIPEEVVEEM
VPEEVVEEVVPEELVEEVIPEEVVEKVVPEQLVEKVVPEELVEKVLPEELVEEVVPEELIEEVVTEEVIEEIIPEEIIEEIIPEEVVEERVSIEVVEELVPEELIEEVVAEEIFEEVVPEKLIEEVV
AEELIEEMVPEVVEEVVEEIIPEEVIEEVIPEEIREEIIPEEVVEEVIEEVIPEELIEEEISREIVEEVIPEEVVEEMMPEEVIEEVIPEEIREEMIPEEVVEEMIPEIEIEQVLPEELIEELVPEVEIE
EILPEELIEELVPEGVIEEVIPKELIEEVIPEELIKEVVPEEVVEEVIPKEIVEEVVPEELIEEVIPEELIEEVVPEEMIEEVIPEELIEEVVPEEMIEEVIPEELIEEVVPEEVKEEVIPVELIEEVVP
EEMIEEVISEEVVEEVIPEELVEEVLPEVFVEEIVPKELVEEVIPEEIVEEVIPEELVEEVIPEELVEEVISEELLEEVIPELLVEEVVPEVIEEVVPEVVEEVIPEVVEEVVTEELVEEVIPEEVV
EEMIPGVEIEEVLPEELVEEVIPEEIVEEVISEEDFEEVIPEALVEEVIPELLVEEVVLEELIEEVIPEVVEEVIPEEVVEEVIPEEVVEKVVPEELVEEVIPEELVEEVIPEEVPEEVVPEELVEE
VISEELVEEVIPEEIVEEVIPEEIVEEVMPEEVAEEVVPEELVEEEVPEELVEEVIPEVLVEEVIPEEIVEEVVPEEVVEVVIPEEFIEEIIPEEVVVEVTPEELIEEMKPEELFEEVIPKELIEEIIP
EELIKEVIPEEIIEEVISKEIVEEVVEEEIIEEVIPEEIVEEVVPEELVEKKVPEEIVEEVIYEEFLEEIMPEELIEEMKPEELIEEIIPREIVEEIITEEVVEEMKPEELVEEMVPEEVVEEMVPEEV
VEEMVPEEVVEEVITEKLIEEVVPEEIIEELIPEELVEEVIPEKLIEEMIPEEVVEEMLLEEVVEEVVPEVEIEEMIPEQLVEEVVPEELVEKEIPEVVEEIITEEVVEEMVPEEVVEEMVPEEV
VEEVILEKLIEEVVPVEVIEEVVPEEVIEEVIPEKLIEEIIPEVLVEEVVPEELIEEVIPEVLFEEVVPEVVEEVVPEELVEKVVPEEIVEEVIPEFVEEVISEEIVEEVVTEELIEEMIPEEVVEEVI
PEELVEEVVPVELIEEVVPVELLEEVIPEELVEEVVPVELIEEVVPVELLEEVIPEEVVEEVIPEVVKEVVPEVVEEVVPEVVEEVVPEVVEEVVPEVVEEVVPEELIEEVIPEELIKEVLPEE
VIEELKPEELIEEIIPEEVVEEVIPEELIEEMIPELLVEEEVPEELVEEVIPEVLVEEMVPEEIVEEVIPEELIAEFAPEELFEKVIPEEVVEELVSEELVEEVLPEVVEEVLPEVFEEVIPEEVIEE
IVPEEVMEEVVPKELIEEVIPEELIEEVIPEEVAEEVVPEKLIEEVIPEELIKEVVPEELIEEVITEEVVEEAISEELVEEVVPEEVVEEVIPEELIEEVVPEELIEEIIPEELVEEVIPEELVEEVVP
EELVEEVIPEELVEEVVPEELVEKLIPEELVEEVIPELLVEEVVPEEVVEELIPEELVEEVIPEELVEEVIPEEVPEEVIREELVEEVIPEELVEEVIPEEVPEEVVPEELVQEVIPEELVEEVVH
EEVPEEVVPEKLVEEVIPEELFEEVVPEEVPEEVVPEELVEEVISEELVEEVIPEELVEEFIPEELVEEVIPEELVEEVIPEELVEEFIPEELVEEVIPEELVEEVIPEVLVEEEIPEKLVEEVVP
EELVEEVIPEVLVEEVVTEEVPEEVVPEELVAEVVPEELVKEVIPEELVEEVIPEEIVEEVIPEEVVEEVVPEELVEEVIPEEVVEEVIPEELVEEVVPVELVEEVVPEELVEEVIPEELVEEVI
PEEIVEEVIPEEVVEEVIPEEIVEEVISEELIEEVVPVELLEEVVPVEVLEDVIPEEVVEEVIPEEFIEEMIPEEIIEEAIPEEIVEEVIPEVVEEVVPEELVEEVIPEEVVEKVVPEELVEEVIPEEL
VEEVIPEEVPEEVVPEELVEEVISEELVEEIIPEELVEEVIPEEVVEEVVPEELVEEVIPEELVEEVIPEELVEEVIPEVVEEVIPEKLVEEVVPEELIEEMIPEEIVEEVVPEVVEEVIPEELIEE
VIPEELVEEVIPEKLIEEMIPEELVEEVILEEVVEEVIPEEVVEEVLPEELVEEVVPEELVEEVAPVELLEEVIPEELLEEVIPEELVEEVIPEELVEDVIPEELVEEVIPEELVEEIIPEELVEEVI
PEVLVEEEVPEELVEEVIPEELVEEVIPEELVEELIPEVLEEVIPEKLVEEVVPVELLEEVIPEELIEEVVPEVLVEVIPEKVVEEVIHEEIVKEVVPEELVEEVVPEEIVEEVVPEEVLEEVIPK
VLLEEEIPEKLVEEVIPEELIEEVVPEELVEEVMPEEVVEEVVPEELVEEVIPEEVVEEVIPEELVEEVVPVELLEEIVPVELLEEVIPEELVEEVIPEELVEEVVPEELVAEVIPEKLVEEVIPE
KLVEEVIPEELVEEVVPEELVAEVIPEKLVEEVIPEELVEEIIPEELVEEVVPEELVEEVVPEELVEEVIPEELVEEVIPEVLVEEEVPEELVEEVIPEELVEEMLPEELIEVVVPEELVEEVVP
EELVEEVVPEEVVEEVIPEELIEELVPEVEIEEILPEELIEELVPEEIIEEVIPKELIEEVIPEVVVEEVVPEVLVEEVVPEELVEEVIPEVLVEEVPEELVEDVIPEELVEEVVPEELVEEVIPEE
VVEEVLPEELVEEVVPEEIVEEVIPEELVEEEVPEELVEEVIPEEVVEEVIPEEVPEKVIPEKLVEELIPEEVVEELIPEEVVEEMVPIEVVEEVVPTEVLEEVIPEIVVEEVVPEELVEEVIPEE
LVEEVIPEIVVEEVVPEELVEEVIPEELVEEVVPEELVEEVVPEEIVEELLPKEIVEEVIYEEQVEEVIHEELVAEVIPEEVVEEVVPEVVEEVVPEIVAEVIPEEVVEEVVPEVVEEVVPEVVE
EVIPEEVVEEVIPEEFIEEMIPEEIVEEVVPEVVEEVIPEELVAEVIPEELVEEVIPEVLFEEEIPEVVVEEVVPEVVEEEIPEVVVEEVVPEVVEEVIPEEVVEELIPEEVVEEMVPIEIVEEVV
PTEVLEEVIPEIVVEEVVPEELVEEVIPEELVEEVVPEELVEEVVPEVVEEVVPEVVEEVVPEELVEEVIPEVVVEEVVPEELVEEVIPEELVEEIIPEEIVEEVIPEELIEEVVPEVLVEEVIPE
ELVEEVIPEELFEEVVPEELVAEVIPEEIVEEVIPEELIEEFVPEELVEEEVPEELVEEVIPEVLVEEVIPEEFVEEVIPEEIVEEVIPEEIVEEVIPEEVVEEVVPEIVAEVIPEEVVEEVVPEVV
EEVVPEVIEEVVPEVIEEVVPEVVEEVVPEVVEEVVPEVVEEVVPVEVLEEVIPEEVVEEVIPEEFIEEMIPEEIVEEVVPEVVEDVIPEELIAEVVSEELFEEEIPEELVEVVPVELLEEVIPE
ELVEEVIPEELVEEVVPEELVEEVVPEELVEEVIPEELFEEVVPEELVEEVIPEEIVEEVIPEELIEEFVPEELVEEEVPEELVEEVIPEVLVEEVIPEEFVEEVIPEELVEEVIPEELFEEVVPE
ELVEEVIPEEIVEEVIPEELIEEVVPEVLVEEVIPEELVEEVIPEELFEEVVPEELVAEVIPEEIVEEVIPEELIEEFVPEELVEEEVPEELVEEVIPEVLVEEEVPEELVEEVIPEELVEEMLPEE
LIEVVVPEELVEEVVPEIVEEVVPEELVEEVIPEEVVEEVLPEKLVEEVVPEIVEEVAPEVVEEVIPEELVEEIIPEILVEEVVPIEVVEEAVPEVVVEEVVPEKIVEEVVPEKVLEEVIPELLVE
EIVPIEVVGEIVPEEVVEKIVPMEVVEELLPEVVEEVPHEIVEEEVVKEVIPEDEKKVEIVEIIIEEKVEKIPKKKVIEKKKEKEKIRKPKKKVEKEIKPKEVKGVKGKLKKDVKAKHLKGSKVKI
TEDIQKKMEDDIKEKIMEKELKEEIKDKEYLRKVIPKEEKKEPIPKKGKSVEPKKEKQAGVKKEKVPETKKEKAPAPKKEKVQAPKKEKVPAPKKEKPAVPKKEKVPEPKKEKVPEPKKE
KAPAPKKEKVQAPKKEKAPAPKKEKAPAPKKEKAPAPKKEKAPAPKKEKPAVPKKEKPAVPKKEKVPETKKEKVPETKKEKVPETKKEKVPVPKKEKVPAPKKEKVPAPKKEKVPAPK
KEKPAVPKKEKVPETKKEKAPEPKKEKAPEPKKEKAPEPKKEKAPEPKKEKAPAPKKEKVPVPKKEKVPAPKKEKVPAPKKEKPAVPKKEKVPETKKEKEKDSQPKKEKAKETSRPPK
KEKEKGK

**Fig. S4. Glutamic acid–rich repeats in RESA (A), RESA3 (B), LSA3 (C), Pfs230 (D) and Pf11.1 (E).** Sequences from Uniprot database. “EENVEE” repeats are highlighted in pink, “EEVGEE” in green, “EELVEE” in light blue, “EEVVEE” in dark blue and other repeats following the “EEXXEE” pattern in yellow.

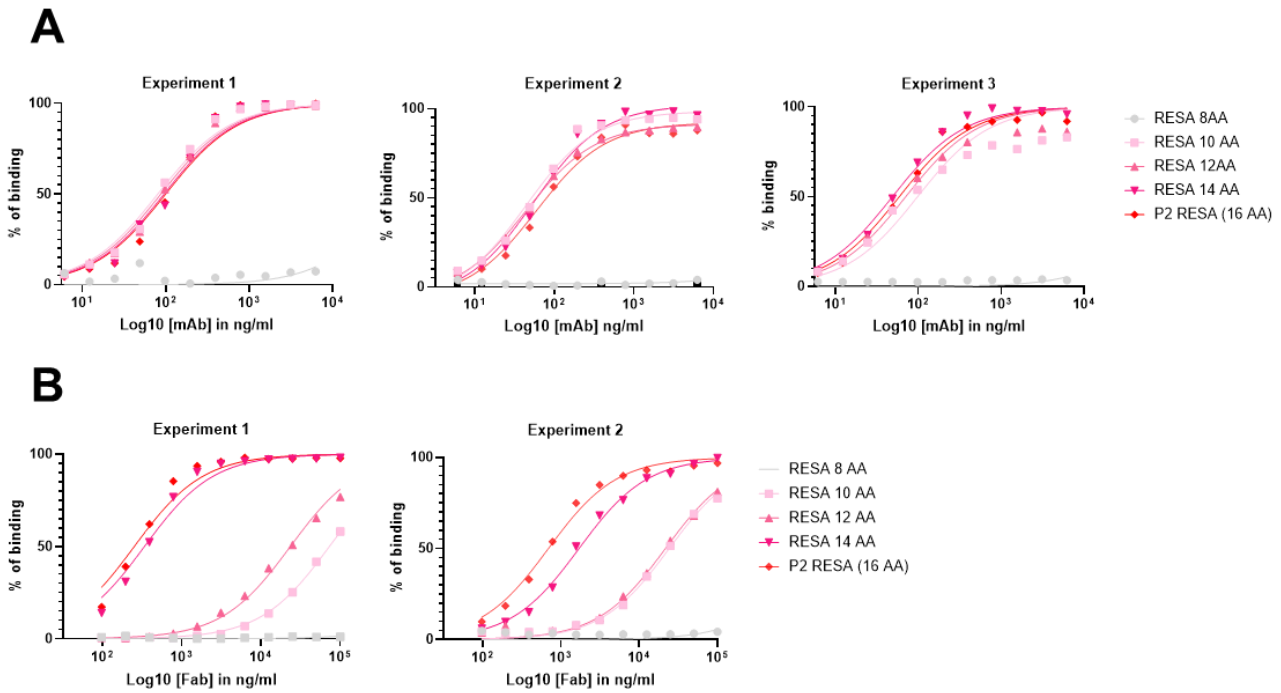


**Fig. S5. BE11K ELISA binding curves to RESA peptides**

Binding to RESA peptides in ELISA: for B1E11K mAb (A), three independent experiments; for B1E11K Fab (B), two independent experiments. Curves were used to calculate EC$_{50}$s shown in figure 4.

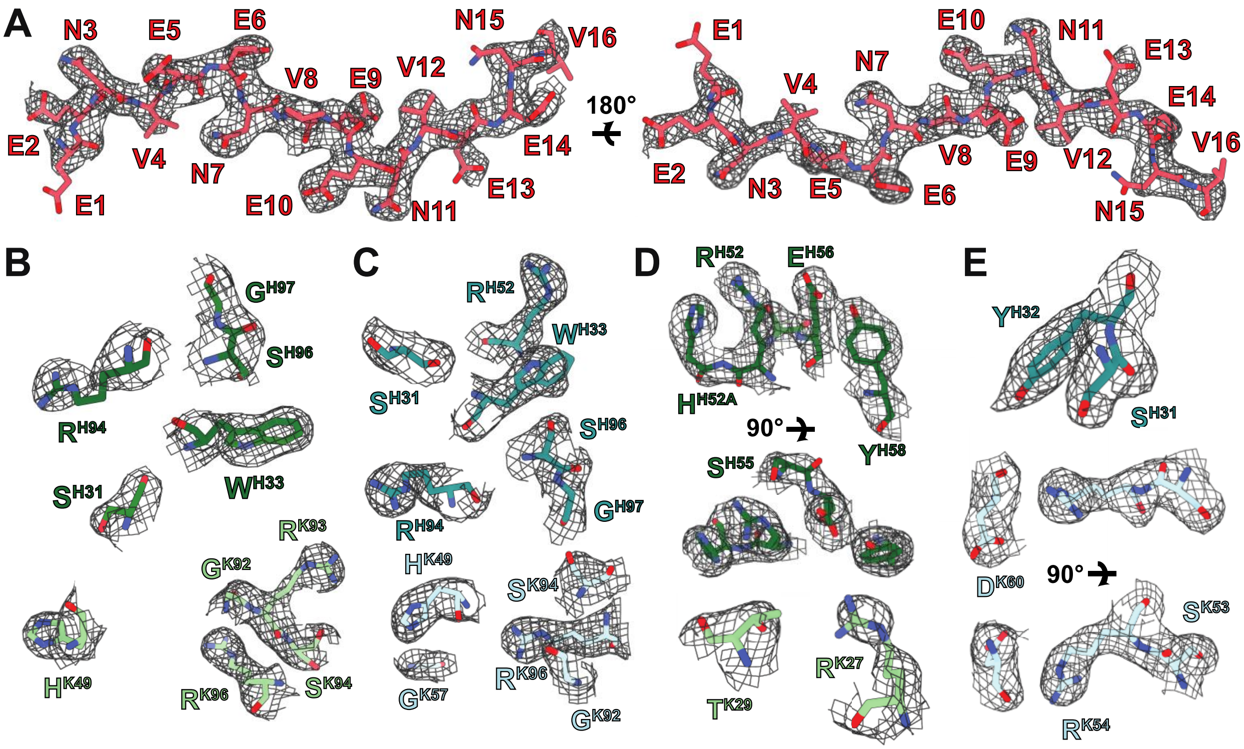


**Fig. S6. Composite omit maps of residues involved in inter-chain interactions.** Composite omit maps of (A) RESA P2 (16AA) peptide (red) which contain the repetitive elements. Composite omit maps of the residues of (B) B1E11K Fab A (heavy chain in green and kappa chain in light green) and (C) B1E11K Fab B (heavy chain in teal and kappa chain in light blue) that interact with the RESA P2 peptide. Composite omit map of residues in (D) B1E11K Fab A and (E) B1E11K Fab B involved in homotypic interaction interface (same coloring scheme).

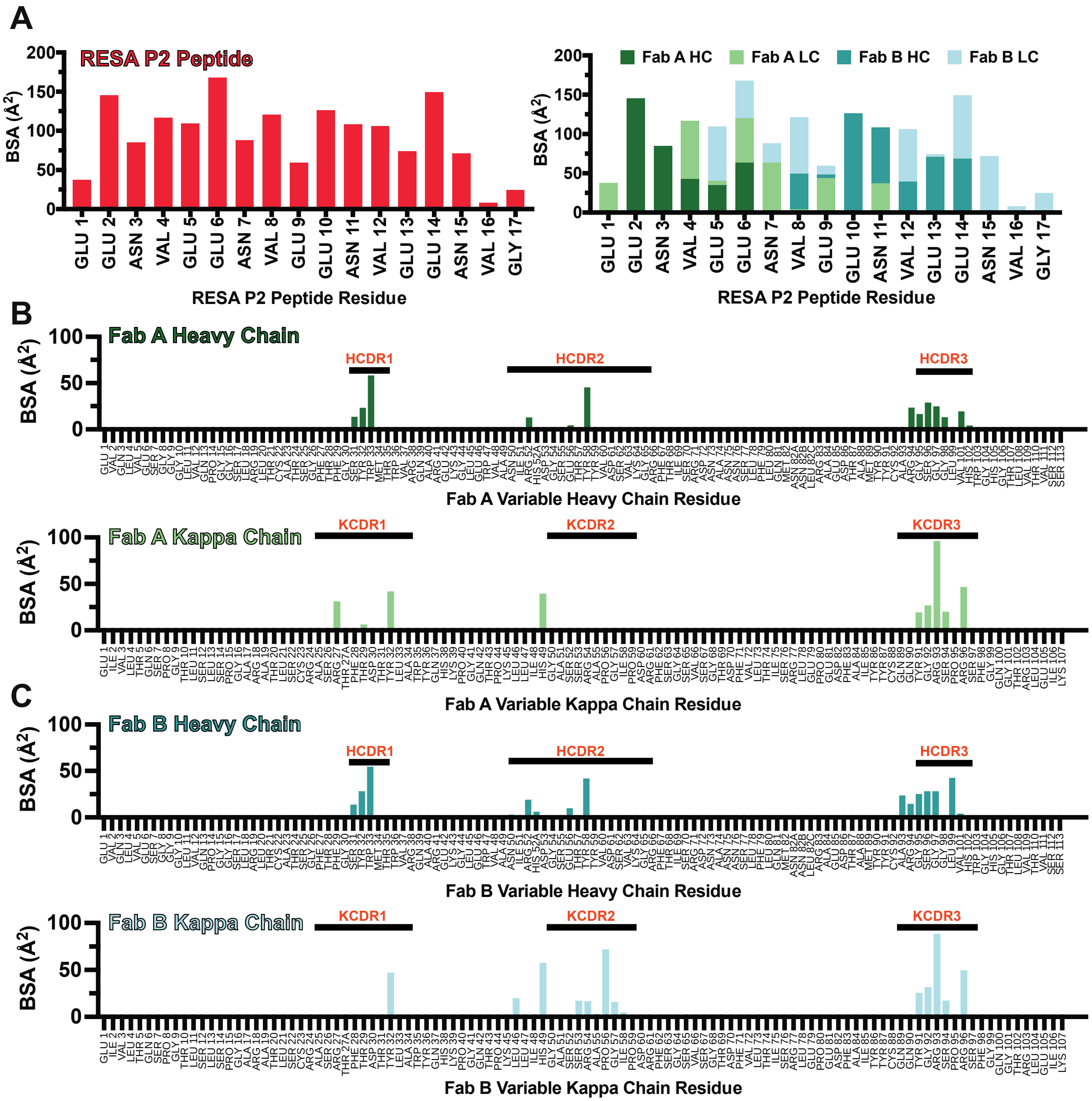


**Fig. S7. Buried surface area plots of B1E11K Fabs and RESA P2 (16AA) peptide interactions.** Bar graphs of the buried surface area of each residue in the (A) RESA P2 (16AA) peptide and both heavy and kappa chains of (B) B1E11K Fab A and (C) B1E11K Fab B. Kabat numbered CDRs are marked with bars.

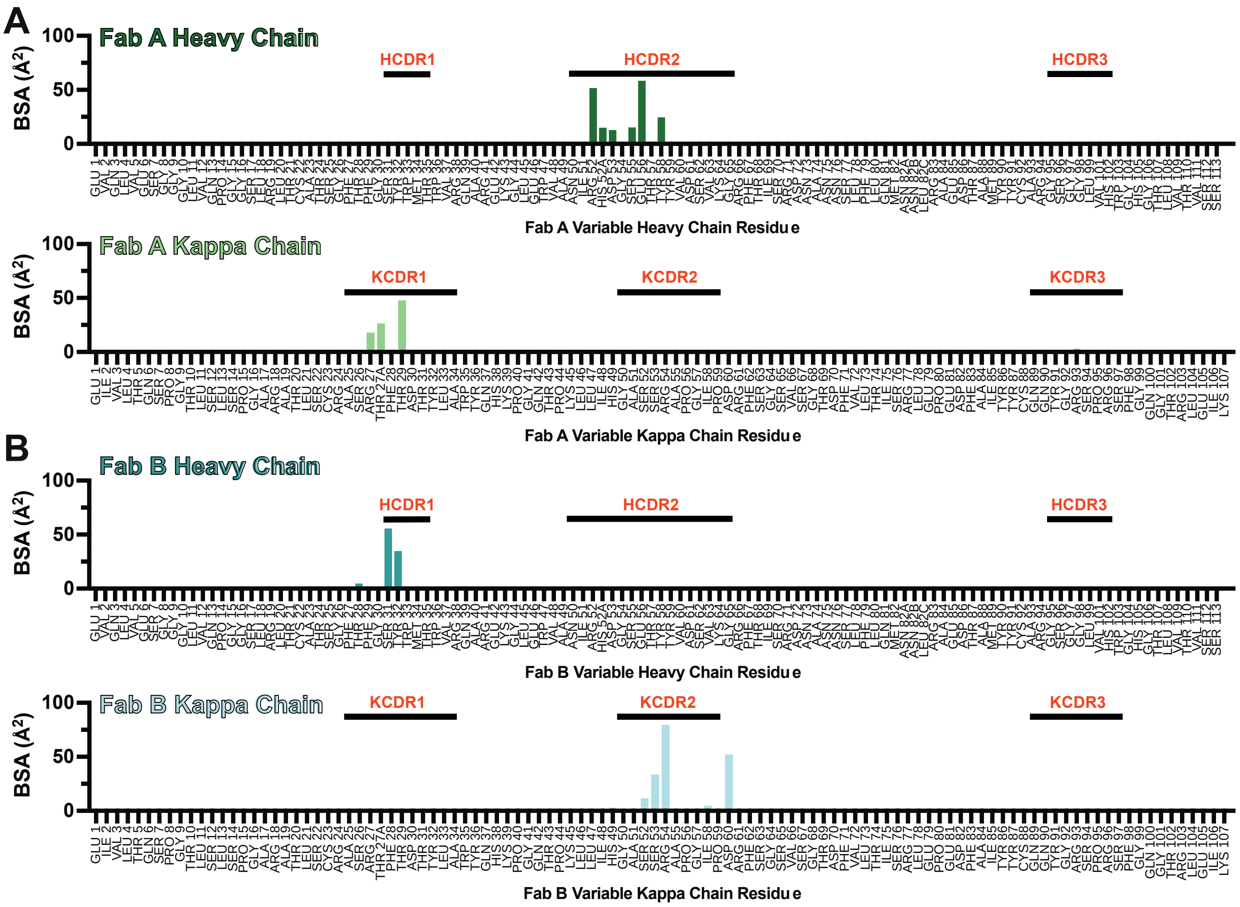


**Fig. S8. Buried surface area plots of B1E11K Fabs of residues buried at the homotypic interaction interface.** Bar graphs of the buried surface area of each residue of both the heavy and kappa chain of (A) B1E11K Fab A and (B) B1E11K Fab B. Kabat numbered CDRs are marked with bars.

**A**

```
                                        <----------------------------------------FR1---------------------------------------->
                                        E  V  Q  L  V  E  S  G  G  G  L  V  Q  P  G  G  S  L  R  L  T  C  A  T  S  G  F  T  F  G
                    B1E11K-HC     1    GAGGTGCAGCTGGTGGAGTCTGGGGGAGGCCTGGTCCAGCCTGGGGGGTCCCTGAGACTCACCTGTGCGACCTCTGGATTCACTTTTGGT  90
V  91.9% (271/295)  IGHV3-7*01    1    ..............................T.............................T.......AG.............C...A..  90
                                        E  V  Q  L  V  E  S  G  G  G  L  V  Q  P  G  G  S  L  R  L  S  C  A  A  S  G  F  T  F  S
V  91.5% (270/295)  IGHV3-7*03    1    ..............................T.............................T.......AG.............C...A..  90
V  91.5% (269/294)  IGHV3-7*02    1    ..............................T.............................T.......AG.............C...A..  90

                                       <----CDR1-----><------------------FR2-----------------><---------------------CDR2------
                                        S  Y  W  M  T  W  V  R  Q  A  R  E  K  G  L  E  W  V  A  N  I  R  H  D  G  S  E  T  Y  Y
                    B1E11K-HC     91   AGTTATTGGATGACCTGGGTCCGCCAGGCTCGAGAGAAGGGGCTGGAGTGGGTGGCCAATATAAGACACGATGGCAGTGAAACATACTAT  180
V  91.9% (271/295)  IGHV3-7*01    91   ..C..........G.................C..G........................C....AG..A.....A.....G.A.......  180
                                        S  Y  W  M  S  W  V  R  Q  A  P  G  K  G  L  E  W  V  A  N  I  K  Q  D  G  S  E  K  Y  Y
V  91.5% (270/295)  IGHV3-7*03    91   ..C..........G.................C..G........................C....AG..A.....A.....G.A.......  180
V  91.5% (269/294)  IGHV3-7*02    91   ..C..........G.................C..G...A....................C....AG..A.....A.....G.A.......  180

                                       -----------------><------------------------------------------------FR3---------------------
                                        V  D  S  V  K  G  R  F  T  I  S  R  D  N  A  N  N  S  L  F  L  Q  M  N  N  L  R  A  E  D
                    B1E11K-HC     181  GTGGACTCTGTGAAGGGCCGATTCACCATCTCCAGAGACAACGCCAACAACTCACTGTTTCTGCAAATGAACAACCTGAGAGCCGAAGAC  270
V  91.9% (271/295)  IGHV3-7*01    181  ...............................................G..........A..............G............G...  270
                                        V  D  S  V  K  G  R  F  T  I  S  R  D  N  A  K  N  S  L  Y  L  Q  M  N  S  L  R  A  E  D
V  91.5% (270/295)  IGHV3-7*03    181  ...............................................G..........A..............G............G...  270
V  91.5% (269/294)  IGHV3-7*02    181  ...............................................G..........A..............G............G...  270

                                       ----------------------><--------CDR3--------><--------------FR4-------------->
                                        T  A  M  Y  Y  C  A  R  G  S  G  G  L  V  H  W  G  H  G  T  L  V  T  V  S  S
                    B1E11K-HC     271  ACGGCTATGTACTACTGTGCGCGAGGTTCGGGGGGTTTGGTCCACTGGGGCCACGGAACCCTGGTCACCGTCTCCTCA  348
V  91.9% (271/295)  IGHV3-7*01    271  ......G....T.........A...-----------------------------------------------------  295
                                        T  A  V  Y  Y  C  A  R
V  91.5% (270/295)  IGHV3-7*03    271  .....CG....T.........A...-----------------------------------------------------  295
V  91.5% (269/294)  IGHV3-7*02    271  ......G....T.........A..------------------------------------------------------  294
```

**B**

```
                                         <--------------------------------FR1--------------------------------><---------------CDR1-
                                        E  I  V  L  T  Q  S  P  G  T  L  S  L  S  S  G  A  R  A  T  L  S  C  R  A  S  R  T  F  T
                    B1E11K-KC     1    GAAATTGTGTTGACGCAGTCTCCAGGCACCCTGTCTTTGTCTTCAGGGGCAAGAGCCACCCTCTCCTGCAGGGCCAGTCGGACTTTCACC  90
V  92.0% (264/287)  IGKV3-20*01   1    ..........................................C......A.............................A..G.G.T.G.  90
                                        E  I  V  L  T  Q  S  P  G  T  L  S  L  S  P  G  E  R  A  T  L  S  C  R  A  S  Q  S  V  S
V  90.2% (259/287)  IGKV3D-20*01  1    .........................C................C......A...................G.........A..G.G.T.G.  90
V  91.5% (249/272)  IGKV3D-20*02  1    .........................C................C......A.............................A..G.G.T.G.  90

                                       --------------><--------------------FR2--------------------><-------CDR2--------><--------
                                        D  T  Y  L  A  W  Y  Q  H  K  P  G  Q  T  P  K  L  L  I  H  G  A  S  S  R  A  P  G  I  P
                    B1E11K-KC     91   GACACCTACTTGGCCTGGTACCAGCACAAACCTGGCCAGACTCCCAAGCTCCTCATCCATGGTGCATCCAGCAGGGCCCCTGGCATCCCA  180
V  92.0% (264/287)  IGKV3-20*01   91   AG..G......A..............G............G......G..........T....................A...........  180
                                        S  S  Y  L  A  W  Y  Q  Q  K  P  G  Q  A  P  R  L  L  I  Y  G  A  S  S  R  A  T  G  I  P
V  90.2% (259/287)  IGKV3D-20*01  91   AG..G......A..............G..........T.G.G....G..........T...A................A...........  180
V  91.5% (249/272)  IGKV3D-20*02  91   AG..G......A..............G............G......G..........T...A................A...........  180

                                       --------------------------------------FR3--------------------------------------------><--
                                        D  R  F  S  G  S  V  S  G  T  D  F  V  L  T  I  S  R  L  E  P  E  D  F  A  I  Y  Y  C  Q
                    B1E11K-KC     181  GACAGGTTCAGTGGCAGTGTGTCTGGGACAGACTTCGTTCTCACCATCAGCAGACTGGAGCCTGAAGATTTTGCAATATATTACTGTCAG  270
V  92.0% (264/287)  IGKV3-20*01   181  ...................G................AC.....................................G.G............  270
                                        D  R  F  S  G  S  G  S  G  T  D  F  T  L  T  I  S  R  L  E  P  E  D  F  A  V  Y  Y  C  Q
V  90.2% (259/287)  IGKV3D-20*01  181  ...................G................AC.....................................G.G............  270
V  91.5% (249/272)  IGKV3D-20*02  181  ...................G................AC.....................................G.C............  270

                                       --------CDR3-----------><------------FR4------------>
                                        Q  Y  G  R  S  P  R  S  F  G  Q  G  T  R  L  E  I  K
                    B1E11K-KC     271  CAATATGGTCGCTCACCGCGCAGCTTTGGCCAGGGGACTAGACTGGAGATCAAA  324
V  92.0% (264/287)  IGKV3-20*01   271  ..G......A.......-------------------------------------  287
                                        Q  Y  G  S  S  P
V  90.2% (259/287)  IGKV3D-20*01  271  ..G......A.......-------------------------------------  287
V  91.5% (249/272)  IGKV3D-20*02  271  ..----------------------------------------------------  272
```

**Fig. S9. IgBLAST of B1E11K heavy chain and light chain.** Amino acid sequence alignments of the B1E11K heavy chain and light chain with the (A) IGHV3-7 and (B) IGKV3-20 loci. Residues that have undergone somatic hypermutation and partake in electrostatic interactions with RESA are highlighted in yellow. Residues that have undergone somatic hypermutation and take part in homotypic interactions (electrostatic) are highlighted in cyan. Residues that partake in both types of interactions are highlighted in green.

**Table S1. List of isolated monoclonal antibodies.** Genetic characteristics of isolated antibodies. Sequences were analyzed using the IMGT database. Lambda-antibodies are listed over a yellow background and kappa-antibodies over a blue background. V region identity is indicated as a percentage of nucleotides. Antibodies that differ from one another solely through their light chain are indicated using dashed lines (B1C5L/K, B1C8L/K and B1D3L/K).

| | Heavy Chain | | | | Light Chain | | | |
|---|---|---|---|---|---|---|---|---|
| mAb | V gene | J gene | CDR3 length | V region identity % | V gene | J gene | CDR3 length | V region identity % |
| B1C5L | IGHV1-8*01 | IGHJ5*02 | 16 | 93.06% | IGLV1-44*01 | IGLJ3*02 | 11 | 95.44% |
| B1C5K | IGHV1-8*01 | IGHJ5*02 | 16 | 93.06% | IGKV1-5*03 | IGKJ2*02 | 9 | 89.73% |
| B2C10L | IGHV1-2*02 | IGHJ5*02 | 11 | 93.09% | IGLV2-11*01 | IGLJ1*01 | 9 | 96.53% |
| B2E9L | IGHV3-33*01 | IGHJ4*02 | 17 | 96.53% | IGLV1-51*02 | IGLJ2*01 | 10 | 95.79% |
| B1C8L | IGHV4-61*02 | IGHJ5*02 | 11 | 80.70% | IGLV1-51*02 | IGLJ2*01 | 11 | 86.67% |
| B1C8K | IGHV4-61*02 | IGHJ5*02 | 11 | 80.70% | IGKV3-15*01 | IGKJ1*01 | 10 | 96.42% |
| B1D3L | IGHV1-18*01 | IGHJ6*03 F | 21 | 91.67% | IGLV1-47*01 | IGLJ3*02 | 10 | 88.41% |
| B1D3K | IGHV1-18*01 | IGHJ6*03 F | 21 | 91.67% | IGKV2D-29*01 | IGKJ1*01 | 9 | 97.62% |
| B1F9K | IGHV4-4*07 | IGHJ2*01 | 14 | 58.22% | IGKV1-12*01 | IGKJ3*01 | 9 | 100% |
| B1E7K | IGHV3-33*01 | IGHJ4*02 | 17 | 95.14% | IGKV3-11*01 | IGKJ2*04 | 10 | 97.47% |
| B1C3L | IGHV4-4*07 | IGHJ5*02 | 17 | 100.00% | IGLV2-14*01 | IGLJ2*01 | 10 | 90% |
| B2D10L | IGHV4-34*03 | IGHJ5*02 | 22 | 85.61% | IGLV3-1*01 | IGLJ2*01 | 10 | 96.06% |
| B2F7L | IGHV3-74*03 | IGHJ4*02 | 19 | 85.42% | IGLV2-8*01 | IGLJ1*01 | 10 | 100% |
| B1E11K | IGHV3-7*01 | IGHJ4*01 | 9 | 90.74% | IGKV3-20*01 | IGKJ2*04 | 9 | 90.20% |

**Table S2. Raw oocyst count data from four independent standard membrane feeding assays (exp #1 - #4). These data were used to calculate transmission reducing activity shown in Figure 2b. FCS was used as a negative control to calculate TRA, while 2A2 was included as a complement-dependent positive control mAb.**

| exp | sample | conc (μg/mL) | oocyst counts per mosquito | | | | | | | | | | | | | | | | | | | | | Mean |
|---|---|---|---|---|---|---|---|---|---|---|---|---|---|---|---|---|---|---|---|---|---|---|---|---|
| | FCS | | 8 | 0 | 2 | 12 | 0 | 18 | 3 | 9 | 0 | 19 | 0 | 0 | 23 | 4 | 6 | 9 | 0 | 0 | 0 | 1 | | 5.7 |
| | FCS | | 23 | 18 | 28 | 12 | 2 | 12 | 13 | 3 | 9 | 0 | 8 | 10 | 29 | 11 | 4 | 0 | 22 | 16 | 9 | 3 | | 11.6 |
| | 2A2 | 10 | 1 | 0 | 0 | 0 | 0 | 0 | 0 | 0 | 0 | 0 | 0 | 1 | 1 | 0 | 0 | 0 | 0 | 0 | 0 | 0 | | 0.2 |
| | VRC01 | 500 | 14 | 6 | 2 | 0 | 8 | 11 | 4 | 18 | 9 | 17 | 23 | 15 | 12 | 14 | 10 | 16 | 7 | 3 | 4 | 8 | | 10.1 |
| | B1D3L | 500 | 0 | 0 | 0 | 0 | 0 | 0 | 0 | 0 | 1 | 0 | 3 | 0 | 0 | 0 | 2 | 0 | 2 | 0 | 2 | 0 | | 0.5 |
| | B1E7K | 500 | 12 | 0 | 13 | 5 | 15 | 21 | 9 | 17 | 7 | 1 | 19 | 16 | 10 | 2 | 6 | 0 | 5 | 16 | 4 | 7 | | 9.3 |
| #1 | B1C3L | 500 | 7 | 22 | 9 | 0 | 11 | 5 | 0 | 8 | 23 | 1 | 9 | 4 | 2 | 0 | 13 | 2 | 0 | 7 | 6 | 10 | | 7.0 |
| | B1C8L | 500 | 1 | 0 | 0 | 0 | 0 | 0 | 0 | 0 | 0 | 0 | 0 | 1 | 1 | 0 | 0 | 0 | 2 | 0 | 0 | 0 | | 0.3 |
| | B1F9K | 500 | 0 | 2 | 0 | 0 | 2 | 1 | 0 | 1 | 2 | 0 | 1 | 0 | 0 | 0 | 2 | 0 | 3 | 0 | 0 | 0 | | 0.7 |
| | B1C5L | 500 | 2 | 0 | 0 | 0 | 1 | 0 | 1 | 0 | 0 | 0 | 1 | 0 | 1 | 3 | 0 | 2 | 0 | 0 | 1 | 4 | | 0.8 |
| | B1C5K | 500 | 0 | 0 | 0 | 0 | 1 | 1 | 0 | 0 | 1 | 0 | 0 | 0 | 1 | 0 | 2 | 3 | 0 | 0 | 2 | 0 | | 0.6 |
| | B2C10L | 500 | 0 | 2 | 0 | 1 | 1 | 0 | 0 | 0 | 1 | 2 | 0 | 0 | 1 | 0 | 0 | 0 | 1 | 0 | 0 | 0 | | 0.5 |
| | B2E9L | 500 | 0 | 0 | 0 | 0 | 0 | 0 | 6 | 0 | 4 | 0 | 0 | 0 | 2 | 1 | 3 | 0 | 0 | 1 | 0 | 4 | | 1.1 |
| | FCS | | 1 | 12 | 2 | 0 | 3 | 7 | 8 | 3 | 8 | 2 | 6 | 2 | 3 | 3 | 10 | 6 | 5 | 9 | 6 | 12 | | 5.4 |
| | FCS | | 3 | 3 | 2 | 6 | 4 | 7 | 3 | 9 | 4 | 7 | 2 | 1 | 0 | 4 | 8 | 11 | 3 | 15 | 19 | 2 | | 5.7 |
| | 2A2 | 10 | 0 | 0 | 0 | 0 | 0 | 0 | 0 | 0 | 0 | 0 | 0 | 0 | 0 | 0 | 0 | 0 | 0 | 0 | 0 | 0 | | 0.0 |
| | B1D3L | 500 | 0 | 0 | 0 | 0 | 0 | 0 | 0 | 0 | 0 | 3 | 0 | 0 | 0 | 0 | 0 | 0 | 0 | 0 | 0 | 0 | | 0.2 |
| #2 | B1C8L | 500 | 0 | 0 | 0 | 0 | 0 | 0 | 2 | 2 | 0 | 0 | 1 | 0 | 0 | 0 | 1 | 0 | 0 | 0 | 0 | 0 | | 0.3 |
| | B2E9L | 500 | 1 | 0 | 2 | 0 | 0 | 0 | 0 | 0 | 0 | 0 | 0 | 0 | 0 | 1 | 0 | 0 | 0 | 0 | 0 | 0 | | 0.2 |
| | B1F9K | 500 | 0 | 0 | 0 | 0 | 12 | 3 | 0 | 1 | 0 | 1 | 0 | 0 | 0 | 1 | 0 | 0 | 0 | 0 | 0 | 0 | | 0.9 |
| | B1C5L | 500 | 2 | 0 | 1 | 0 | 0 | 0 | 0 | 0 | 0 | 0 | 2 | 0 | 0 | 0 | 4 | 0 | 0 | 0 | 1 | 0 | | 0.5 |
| | B1C5K | 500 | 0 | 0 | 2 | 0 | 0 | 0 | 0 | 0 | 0 | 0 | 2 | 0 | 0 | 0 | 0 | 0 | 2 | 0 | 1 | 0 | | 0.4 |
| | B2C10L | 500 | 0 | 0 | 0 | 0 | 0 | 0 | 0 | 0 | 0 | 0 | 0 | 0 | 0 | 0 | 0 | 0 | 0 | 0 | 0 | 0 | | 0.0 |
| | FCS | | 2 | 6 | 3 | 11 | 15 | 3 | 7 | 19 | 0 | 12 | 2 | 5 | 7 | 2 | 6 | 1 | 6 | 9 | 7 | 5 | | 6.4 |
| | FCS | | 19 | 18 | 11 | 6 | 2 | 8 | 11 | 21 | 0 | 5 | 0 | 5 | 17 | 27 | 13 | 5 | 5 | 13 | 18 | 7 | | 10.6 |
| | 2A2 | 10 | 0 | 0 | 0 | 0 | 0 | 0 | 0 | 0 | 0 | 0 | 0 | 0 | 0 | 0 | 0 | 0 | 0 | 0 | 0 | 0 | | 0.0 |
| | VRC01 | 500 | 5 | 1 | 2 | 0 | 15 | 2 | 2 | 4 | 4 | 6 | 1 | 7 | 7 | 10 | 13 | 7 | 8 | 27 | 7 | 9 | | 6.9 |
| | B1E11k | 500 | 10 | 6 | 13 | 18 | 12 | 7 | 14 | 9 | 10 | 11 | 9 | 6 | 8 | 0 | 9 | 13 | 8 | 7 | 17 | 12 | | 10.0 |
| | B2D10L | 500 | 15 | 14 | 21 | 12 | 28 | 21 | 18 | 19 | 30 | 1 | 2 | 4 | 17 | 12 | 6 | 9 | 10 | 8 | 12 | 8 | | 13.4 |
| | B2F7L | 500 | 9 | 12 | 10 | 6 | 13 | 12 | 8 | 15 | 17 | 16 | 32 | 11 | 4 | 24 | 14 | 5 | 4 | 18 | 9 | 14 | | 12.7 |
| #3 | B1C8K | 500 | 6 | 4 | 5 | 6 | 8 | 8 | 9 | 6 | 17 | 3 | 5 | 21 | 6 | 7 | 14 | 6 | 6 | 18 | 7 | 2 | | 8.2 |
| | VRC01 | 100 | 19 | 11 | 8 | 5 | 18 | 8 | 21 | 16 | 14 | 18 | 21 | 6 | 22 | 9 | 10 | 1 | 4 | 11 | 4 | 13 | | 12.0 |
| | B1C5K | 100 | 8 | 14 | 9 | 16 | 7 | 8 | 9 | 10 | 12 | 24 | 6 | 3 | 17 | 19 | 14 | 9 | 15 | 18 | 9 | 13 | | 12.0 |
| | B1C5L | 100 | 3 | 18 | 3 | 4 | 18 | 21 | 0 | 23 | 14 | 5 | 14 | 13 | 9 | 13 | 4 | 12 | 5 | 17 | 6 | 18 | | 11.0 |
| | B1D3L | 100 | 11 | 3 | 8 | 18 | 6 | 0 | 13 | 19 | 6 | 13 | 5 | 5 | 12 | 0 | 8 | | | | | | | 8.5 |
| | B2C10L | 100 | 3 | 0 | 6 | 5 | 2 | 4 | 0 | 0 | 0 | 4 | 7 | 0 | 9 | 1 | 0 | 3 | 0 | 0 | 3 | 2 | | 2.5 |
| | B2E9L | 100 | 9 | 1 | 2 | 5 | 22 | 6 | 16 | 8 | 17 | 19 | 11 | 17 | 1 | 8 | 4 | 11 | 7 | 1 | 12 | 5 | | 9.1 |
| | B1C8L | 100 | 3 | 9 | 3 | 6 | 11 | 10 | 4 | 3 | 4 | 9 | 9 | 8 | 1 | 8 | 0 | 2 | 6 | 3 | 6 | 10 | | 5.8 |
| | B1F9K | 100 | 7 | 14 | 9 | 12 | 13 | 21 | 8 | 3 | 6 | 8 | 6 | 4 | 13 | 11 | 8 | 19 | 6 | 21 | 18 | 10 | | 10.9 |
| | FCS | | 19 | 15 | 18 | 43 | 12 | 5 | 24 | 35 | 3 | 9 | 14 | 11 | 10 | 27 | 32 | 10 | 5 | 19 | 24 | 19 | | 17.7 |
| | FCS | | 16 | 11 | 30 | 34 | 9 | 72 | 21 | 22 | 22 | 16 | 16 | 29 | 35 | 34 | 28 | 8 | 6 | 18 | 24 | 36 | | 24.4 |
| | 2A2 | 10 | 0 | 0 | 0 | 0 | 0 | 0 | 0 | 0 | 0 | 0 | 0 | 0 | 0 | 0 | 0 | 0 | 0 | 0 | 0 | 0 | | 0.0 |
| | VRC01 | 100 | 30 | 28 | 26 | 5 | 8 | 15 | 19 | 33 | 25 | 27 | 35 | 28 | 33 | 45 | 41 | 14 | 11 | 19 | 11 | 52 | | 25.3 |
| | B1D3L | 100 | 8 | 1 | 8 | 4 | 6 | 16 | 1 | 7 | 0 | 2 | 9 | 0 | 1 | 2 | 8 | 6 | 7 | 19 | 3 | 5 | | 5.7 |
| #4 | B1C8L | 100 | 4 | 2 | 4 | 0 | 0 | 1 | 3 | 1 | 4 | 0 | 0 | 0 | 0 | 0 | 4 | 0 | 4 | 0 | 0 | 3 | | 1.5 |
| | B1F9K | 100 | 3 | 2 | 9 | 6 | 0 | 2 | 13 | 5 | 3 | 1 | 2 | 4 | 0 | 5 | 4 | 0 | 12 | 13 | 13 | 2 | | 5.0 |
| | B1C5L | 100 | 0 | 6 | 0 | 11 | 9 | 0 | 8 | 1 | 18 | 6 | 0 | 2 | 9 | 4 | 3 | 1 | 2 | 3 | 18 | 2 | | 5.2 |
| | B1C5K | 100 | 6 | 0 | 2 | 18 | 1 | 1 | 7 | 0 | 0 | 0 | 0 | 1 | 18 | 1 | 3 | 3 | 14 | 1 | 15 | 2 | | 4.7 |
| | B2C10L | 100 | 7 | 0 | 7 | 0 | 0 | 6 | 0 | 0 | 0 | 1 | 9 | 1 | 3 | 1 | 6 | 6 | 5 | 5 | 2 | 1 | | 3.0 |
| | B2E9L | 100 | 2 | 3 | 6 | 2 | 0 | 4 | 2 | 11 | 1 | 10 | 11 | 5 | 3 | 6 | 22 | 1 | 5 | 14 | 18 | 4 | | 6.5 |

**Table S3. Electrostatic Interactions between B1E11K Fabs and RESA P2 (16AA) peptide.**

| RESA P2 | Fab A Heavy | Fab A Kappa | Fab B Heavy | Fab B Kappa |
|---|---|---|---|---|
| E1* | | H49* | | |
| E2* | R94* | | | |
| E2 [O] | S96 [N] | | | |
| E2 [O] | G97 [N] | | | |
| N3 [Nδ2] | S31 [O] | | | |
| N3 [Oδ1] | W33 [N] | | | |
| V4 [N] | S96 [Oγ] | | | |
| V4 [O] | S96 [Oγ] | | | |
| V4 [O] | | R96 [Nη] | | |
| E5 [O] | | R96 [Nη] | | |
| E5 [Oε] | | | | G57 [N] |
| E6* | | R96* | | |
| E6 [Oε] | | S94 [N] | | |
| E6 [O] | | | | H49 [Nε2] |
| N7 [N] | | G92 [O] | | |
| N7 [Nδ2] | | G92 [O] | | |
| E9* | | R93* | | |
| E10* | | | R94* | |
| E10 [O] | | | S96 [N] | |
| E10 [O] | | | G97 [N] | |
| N11 [Nδ2] | | | S31 [O] | |
| N11[Oδ1] | | | W33 [N] | |
| V12 [N] | | | S96 [Oγ] | |
| V12 [O] | | | S96 [Oγ] | |
| V12 [O] | | | | R96 [Nη] |
| E13* | | | R52* | |
| E13 [Oε] | | | W33 [Nε1] | |
| E14* | | | | R96* |
| E14 [Oε] | | | | S94 [N] |
| E14 [Oε] | | | | S94 [Oγ] |
| N15 [N] | | | | G92 [O] |
| N15 [Nδ2] | | | | G92 [O] |

The asterisk (*) denotes a salt bridge. Atoms involved in hydrogen bonding interactions are shown in square brackets.

**Table S4. Electrostatic Homotypic Interactions between B1E11K Fab A and Fab B**

| Fab A Heavy | Fab A Kappa | Fab B Heavy | Fab B Kappa |
|---|---|---|---|
| R52* | | | D60* (FR) |
| H52A* | | | D60* (FR) |
| S55 [Oγ] | | | R54 [Nη1] |
| E56 [Oε] | | | R54 [N] |
| E56* | | | R54* |
| Y58 [Oη] | | | S53 [Oγ] |
| | T29 [O] | Y32 [Oη] | |
| | R27 [Nη] | S31 [O] | |

The asterisk (*) denotes a salt bridge. Atoms involved in hydrogen bonding interactions are shown in square brackets.
